# *D*-*ꞵ*-hydroxybutyrate stabilizes hippocampal CA3-CA1 circuit during acute insulin resistance

**DOI:** 10.1101/2023.08.23.554428

**Authors:** Bartosz Kula, Botond Antal, Corey Weistuch, Florian Gackière, Alexander Barre, Victor Velado, Jeffrey M Hubbard, Maria Kukley, Lilianne R Mujica-Parodi, Nathan A Smith

## Abstract

1.

The brain primarily relies on glycolysis for mitochondrial respiration but switches to alternative fuels such as ketone bodies (KBs) when less glucose is available. Neuronal KB uptake, which does not rely on glucose transporter 4 (GLUT4) or insulin, has shown promising clinical applicability in alleviating the neurological and cognitive effects of disorders with hypometabolic components. However, the specific mechanisms by which such interventions affect neuronal functions are poorly understood. In this study, we pharmacologically blocked GLUT4 to investigate the effects of exogenous KB D-ꞵ-hydroxybutyrate (D-ꞵHb) on mouse brain metabolism during acute insulin resistance (AIR). We found that both AIR and D-ꞵHb had distinct impacts across neuronal compartments: AIR decreased synaptic activity and long-term potentiation (LTP) and impaired axonal conduction, synchronization, and action potential (AP) properties, while D-ꞵHb rescued neuronal functions associated with axonal conduction, synchronization, and LTP.

**Significance statement:** This study investigates the impact of acute insulin resistance on the functionality of the hippocampal circuit and the potential protective effects of ketone body supplementation. By inhibiting GLUT4 receptors to induce acute insulin resistance, we reveal several detrimental changes caused by impaired neuronal glucose uptake. These changes include impairments in synaptic activity, axonal conduction, and neuronal firing properties. The study further examines the distinctive effects of acute insulin resistance and the rescue agent D-βHb on synaptic activity, long-term potentiation, axonal conduction, synchronization, and neuronal firing. By shedding light on neuronal responses during insulin resistance, this investigation advances our understanding of neurological disorders associated with hypometabolism and highlights the potential therapeutic value of D-βHb.

## 3. Introduction

Over the past two decades, research on insulin signaling in the brain has become increasingly important in studies of type 2 diabetes mellitus (T2DM) and Alzheimer’s disease (AD) [1,2]. Insulin plays a critical role in memory formation processes in the hippocampus, and systemic insulin resistance can interfere with hippocampal metabolism and cognitive function, as shown by various studies [3,4]. Additionally, insulin is an essential element in memory processing in the hippocampus and a vital mediator of cognitive impairment in patients with T2DM and AD due to its function in promoting glucose transporter 4 (GLUT4) translocation [5,6]. Although insulin from the blood crosses the blood‒brain barrier, research suggests supplementary local insulin synthesis and release in the brain [7], which enhances the pro-cognitive effects of brain glucose [8]. Therefore, impaired glucose uptake in the hippocampus may contribute to cognitive decline associated with aging, T2DM, and AD [9,10,11].

This relationship has led to the hypothesis that brain hypometabolism reflects encroaching neuronal insulin resistance [12]. The hippocampus is highly enriched with GLUT4 receptors, which serve as the primary neuronal insulin-dependent glucose transporter, especially in brain areas with high concentrations of insulin receptors and relatively high neuronal activity [8,13,14]. Insulin acts by enhancing GLUT4 translocation across neuronal membranes under the regulation of phosphatidylinositol 3-kinase (PI3K), which promotes memory formation [15,16]. During memory formation, GLUT4 is primarily expressed in the soma rather than in the more metabolically active hippocampal neuropil [17,18], suggesting that the primary effect of insulin may be the maintenance of neuronal firing. Mouse models of AD or diet-induced obesity (DIO) showed blunting of the pro-cognitive effects of intrahippocampal insulin injections and decreased local glucose metabolism [19,20], coupled with neuronal hyperexcitability with epileptiform spikes and impairment of GLUT4 translocation [21].

During periods with lower glucose availability, ketone bodies (KBs) supplement metabolism as an alternative fuel source [22,23]. The ketogenic diet has shown pro-cognitive benefits in patients suffering from T2DM or AD. These benefits of the ketogenic diet may arise due to reduced neuronal firing rates during ketosis [24,25], increased ATP:ADP ratios [26], increased secretion of gamma-aminobutyric acid (GABA) due to substantial formation of acetyl-coenzyme A [27], equilibration of the NAD:NADH ratio, and reduced production of free radicals, which together may help to temper neuronal hyperexcitability and its metabolic consequences [28,29].

In addition, KBs are crucial in protecting mitochondria from acute metabolic stress by preventing mitochondrial permeability transition (mPT) through their effects on intracellular calcium levels [30,31]. Furthermore, KBs can affect neuronal firing by promoting the opening of ATP-sensitive K^+^ channels (K-ATP), thus reducing the cytosolic pool of ATP generated from glycolysis [25,31].

Exogenous KBs, such as *D*-β-hydroxy-butyrate (D-ꞵHb) ester, can increase KB levels in circulation without the severe dietary restrictions of the ketogenic diet [32,33]. Whether exogenous KBs maintain their neuroprotective effects in patients with normal plasma glucose levels remains unclear; however, recent studies have shown that in healthy adults, D-ꞵHb led to increased overall brain activity and stabilized functional networks [34,35].

These observations led us to hypothesize that exogenous KBs could be utilized to rescue neuronal metabolism after cerebral insulin resistance. To test this hypothesis, we established a murine hippocampal model of acute insulin resistance (AIR) through the inhibition of GLUT4 by administration of indinavir [36,37]. We then studied how neuron-specific insulin resistance affected hippocampal neurons in the CA3-CA1 circuit, a model circuit for studying learning and memory. We tested the effects of D-ꞵHb during AIR to determine the therapeutic potential of exogenous KBs in hypoglycemic animals. To evaluate the circuit-wide effects on synaptic and axonal function, we obtained field potential recordings in the hippocampi of the treated mice and patch-clamp recordings in hippocampal slices to assess the in vitro effects of AIR and D-ꞵHb on the electrophysiological properties of CA1 pyramidal neurons and CA1 fast-spiking interneurons (FSIs). Finally, we used computational modeling approaches to relate our electrophysiological results and Na^+^/K^+^ ATPase dysfunction as the potential reason for the detrimental changes observed during AIR [38].

## 4. Results

Various paradigms of synaptic plasticity associated with learning and memory have been identified in the hippocampus. Many studies have investigated long-term potentiation (LTP), long-term depression (LTD), spike-timing-dependent plasticity, and excitatory postsynaptic potential (EPSP)-spike potentiation in hippocampal circuits, making the hippocampus a classic system for studying neuroplasticity. Furthermore, the simple cytoarchitecture of the hippocampus makes it an ideal model system. In this study, we utilized indinavir, a potent GLUT4 receptor blocker, to pharmacologically induce AIR in hippocampal slices. Our investigation focused on the stratum radiatum of CA1, where numerous synapses are formed between Schaffer collaterals (SCOs) and the apical dendrites of pyramidal neurons.

### 4.1 Synaptic activity and LTP, but not fiber volleys (FVs), are adversely affected by AIR and are not reversed by either 0.1 mM or 1 mM D-ꞵHb

To test synaptic transmission within CA1 for a wide range of energetic demands, we measured the circuit’s response under 3 different stimulation paradigms. First, we applied mild physiological stimulation (not imposing high energetic demands [39,40]), consisting of paired stimulation applied at 25 Hz, repeated every 20 s over 30-60 trials [**Fig. 1A-F**].

**Figure 1:**
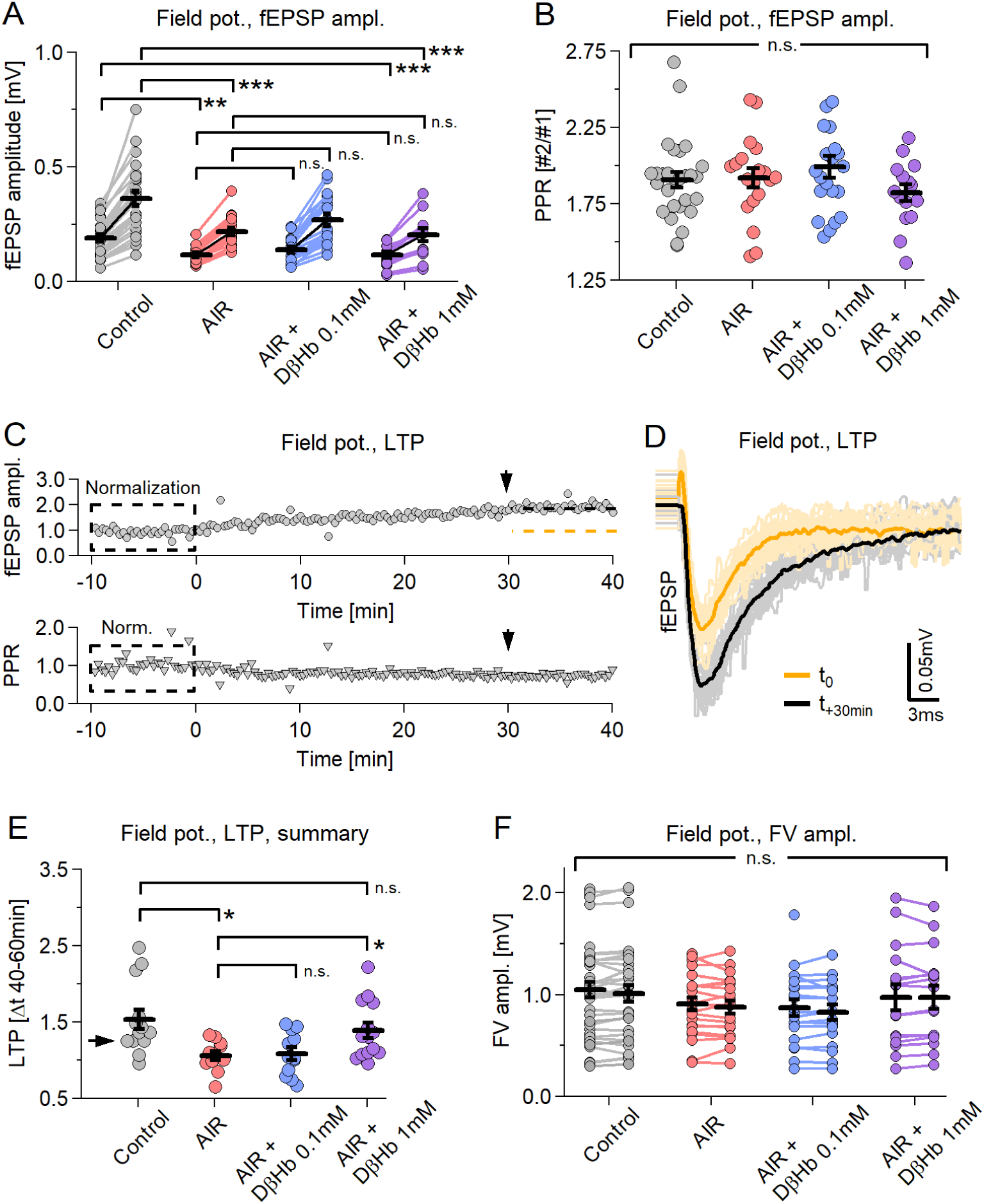
Synaptic activity and LTP, but not neuronal firing or PPR, decrease during paired stimulation under AIR conditions and do not recover during D-ꞵHb administration. A) Scatter plot of the fEPSP amplitudes evoked by paired-pulse stimulation of Schaffer collaterals. Each pair of circles is the average of 30-40 consecutive responses. The black bars represent the mean ± SEM. Control fEPSPs in gray, AIR in red; 0.1 mM D-ꞵHb+AIR in blue; 1 mM D-ꞵHb+AIR in purple. Control: n=26 slices; AIR n=19; 0.1 mM D-ꞵHb+AIR: n=20; 1 mM D-ꞵHb+AIR: n=16. B) Scatter plot of the paired-pulse ratio (PPR) of the fEPSPs recorded in A. The labels, n, are identical to those in A). There were no significant differences (p=0.41, ANOVA). C) On top: Representative recording of a control experiment in which paired stimulation was applied for 50 min. Each circle represents the fEPSP amplitude normalized to the average of the first 30 fEPSPs. The dashed orange line represents the mean amplitude at time −10 to 0 min. The black line represents the mean amplitude at +30 to +40 min. Bottom: corresponding normalized PPR. D) fEPSP waveforms recorded in C), showing LTP development. In gray, 30 fEPSP waveforms at t=0 ± 5 min and their average (black). In yellow, 30 fEPSP waveforms at t=+30 ± 5 min and their average (orange). Stimulation artifacts and FVs are removed. E) Scatter plot of LTP signals triggered by paired stimulation. Control: n=13; AIR n=11; 0.1 mM D-ꞵHb+AIR: n=12; 1 mM D-ꞵHb+AIR: n=14. F) Scatter plot of the FV amplitudes recorded during A). The labels, n, are identical to those in A). There were no significant differences (p=0.42, p=0.40; ANOVA). In all plots, *p<0.05; **p<0.01; ***p<0.001.

During the AIR condition, we observed strong decreases in field excitatory postsynaptic potential (fEPSP) amplitudes, −39.23 ± 8.72% (1st stimulation) and −38.79 ± 8.93% (2nd stimulation), compared to baseline [**Fig. 1A**]. The amplitude did not recover when D-ꞵHb was applied at low (0.1 mM) [**Fig. 1A**, +13.10 ± 6.64% and +14.07 ± 6.76%, p=0.34, p=0.30] or high (1 mM) concentrations [**Fig. 1A**, −5.98 ± 6.13% and −5.81 ± 6.26%, p=0.88; p=0.56]. No changes in the paired-pulse ratio (PPR) were observed in any of the conditions [**Fig. 1B**, Control: 1.89; AIR: 1.92; 0.1 mM D-ꞵHb+AIR: 2.00; 1 mM D-ꞵHb+AIR: 1.88; p=0.54], suggesting that the activity at the synaptic sites remained largely unchanged.

When applied over sufficient time, our paired stimulation triggered LTP in CA1, as evidenced by a gradual increase in the fEPSP amplitude [**Fig. 1C, top**] but no change in the PPR [**Fig. 1C, bottom**]. Therefore, we compared the LTP magnitude between the groups by calculating the ratio of the fEPSP amplitudes during the first 10 min and the amplitudes at 40 ± 10 min [**Fig. 1D-F**]. The control condition showed the strongest LTP ratio of 1.53 ± 0.13. LTP induction was almost abolished during AIR [**Fig. 1E**, 1.06 ± 0.06 ratio; −30.66 ± 9.42% reduction vs. control, p=0.018], and 0.1 mM D-ꞵHb did not reverse this effect [**Fig. 1E**, 1.08 ± 0.08, +1.25 ± 6.61% AIR vs. D-ꞵHb+AIR 0.1 mM, p > 0.99]; however, 1 mM D-ꞵHb did reverse this effect [**Fig. 1E**, 1.39 ± 0.10, +21.19 ± 7.64% AIR vs. D-ꞵHb+AIR 1 mM, p=0.044].

Moreover, we found no significant effects of either AIR or D-ꞵHb+AIR on the amplitudes of the FVs [**Fig. 1F**, Control: 1.03 ± 0.08 mV; AIR: 0.89 ± 0.07; D-ꞵHb+AIR 0.1 mM: 0.85 ± 0.08; D-ꞵHb+AIR 1 mM: 1.03 ± 0.12; p=0.40]. Next, we tested whether the circuit would behave differently under a stronger stimulation that imposed greater metabolic demands. We stimulated the SCO with 20 pulses applied at 25 Hz, with 20 s breaks between trials. To ensure that the onset of LTP did not skew the results, we recorded all train stimulations at least 40 min after the paired stimulation paradigm.

All compared groups showed potentiation of fEPSP amplitudes during the train, reaching a peak at stimuli 5-6. Thereafter, the amplitudes decreased slightly but remained potentiated [**Fig. S1A, B**]. Interestingly, throughout the entire stimulation period, the fEPSP amplitudes matched the differences we observed during the paired stimulation experiments [**Fig. S1B**; at stimulus 5: - 34.25 ± 11.80% Control vs. AIR, p=0.031; −40.54 ± 11.48% Control vs. 1 mM D-ꞵHb+AIR, p=0.000044; +3.37 ± 7.80 AIR vs. 0.1 mM D-ꞵHb+AIR, p = 0.99; −25.21 ± 7.36% AIR vs. 1 mM D-ꞵHb+AIR, p=0.0054]. The FV amplitudes followed a different pattern, with potentiation at stimulus 2, followed by a consistent decline to approximately 70% of the initial amplitude [**Fig. S1C**]. Similar to the paired stimulation experiments, no statistically significant differences were observed between groups at any time point during the train stimulations [**Fig. S1C**; p=0.16 to 0.40].

Finally, we examined the hippocampal circuit under intense nonphysiological stimulation conditions. This involved applying long trains of 50 stimuli at 25 Hz, spanning across 60 trials with a 20 s break between each trial. The responses during this specific stimulation condition varied among trials, making it inappropriate to average the values over the entire stimulation period. As a result, we represent the fEPSP and FV amplitudes as heatmaps. In these heatmaps, each point represents the group’s mean response to a single stimulus at a given time point.

Consistent with our previous results, we observed a period of fEPSP amplitude potentiation followed by a steady decline [**Fig. S2A-D**; at maximum, trial 4-6, stimuli 7-9: −34.81 ± 10.56% Control vs. AIR, p=0.0084; −48.27 ± 11.37% Control vs. 1 mM D-ꞵHb+AIR, p=0.00065; +10.19 ± 11.39% AIR vs. 0.1 mM D-ꞵHb+AIR, p=0.59; −13.66 ± 9.08% AIR vs. 1 mM D-ꞵHb+AIR]. Synaptic depression was first observed during the final stimuli of the stimulation trains, approximately 3 min into the stimulation period, and became more prominent with each subsequent train stimulation application [**Fig. S2A-D**]. In general, the fEPSP amplitudes in the control and AIR groups differed substantially during the first 7 min of the recordings [**Fig. S2A**]. The fEPSP amplitudes in the control and 1 mM D-ꞵHb+AIR groups also differed throughout most of the stimulation trains [**Fig. S2B**], but the fEPSP amplitudes in the 0.1 mM D-ꞵHb+AIR and AIR groups scarcely differed except at the end of the stimulation trains [**Fig. S2C**]. The fEPSPs in the AIR and 0.1 mM D-ꞵHb groups were generally comparable. The FVs remained statistically similar among all the groups, with approximately the same response pattern [**Fig. S2E-G**], replicating the results in Fig. 1 and Fig. S1.

Taken together, these results strongly suggest that the effects of AIR and/or D-ꞵHb could differ among cellular compartments (i.e., dendritic versus somatic) and have different specificity for various cellular compartments, with synapses being particularly susceptible to the metabolic challenges induced by GLUT4 inhibition.

### 4.2 AIR impairs axonal conduction speeds, but D-ꞵHb treatment restores and enhances axonal conduction in a dose-dependent manner

In the previous sections, we highlighted how AIR and D-ꞵHb affected fEPSPs and FVs, finding a surprising lack of adverse effects of AIR on FV amplitudes and a noticeable reduction in fEPSP amplitudes, with the effects further exacerbated by D-βHB. This difference suggests that AIR and/or D-ꞵHb treatment might have differential effects on cellular compartments. We then investigated whether other axonal properties changed during AIR and whether D-βHB application remedied such changes.

We found that inducing AIR resulted in a strong reduction in the conduction velocities (CVs) of SCOs [**Fig. 2C**; −13.07 ± 2.87% Control vs. AIR, p=0.027]. After the addition of 0.1 mM D-ꞵHb during AIR, the CV recovered to control levels [**Fig. 2C**; +12.43 ± 4.42% AIR vs. 0.1 mM D-ꞵHb+AIR, p=0.034; +0.01 ± 0.07% Control vs. 0.1 mM D-ꞵHb+AIR, p=0.99], and the application of 1 mM D-ꞵHb significantly improved the CV, surpassing the control level [**Fig. 2C**; +31.79 ± 4.59% for AIR vs. 1 mM D-ꞵHb+AIR, p=4.2E-8; +19.10 ± 4.53% for control vs. 1 mM D-ꞵHb+AIR, p=0.00057].

**Figure 2:**
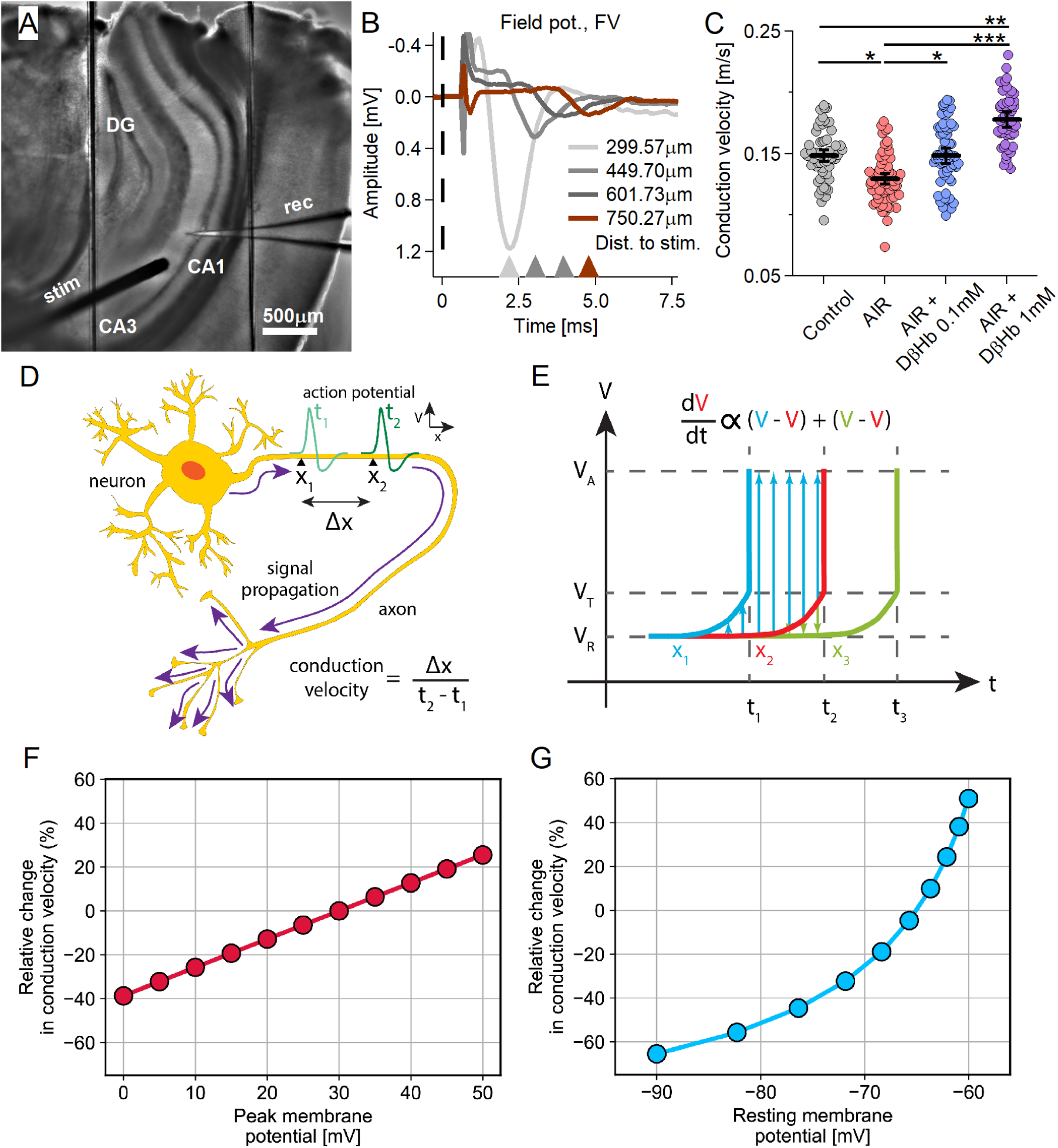
Conduction velocity (CV) of Schaffer collaterals decreases under AIR conditions, is rescued by 0.1 mM D-ꞵHb, and increases with 1 mM D-ꞵHb. A) Representative hippocampal slice during CV recording. B) Representative averaged FV waveforms recorded in CA1. The stimulus onset time (t=0) is marked with a dashed line. Stimulus artifacts are removed. C) CV scatter plots. Each colored circle represents a CV replicate. Control in gray, AIR in light red, 0.1 mM D-ꞵHb+AIR in blue, and 1 mM D-ꞵHb+AIR in purple. Data are shown as the mean ± SEM. *p<0.05; ***p<0.001, nested ANOVA, Control: m=62, n=24, N=14; AIR: m=65, n=22, N=13; 0.1 mM D-ꞵHb+AIR: m=64, n=21, N=13; 1 mM D-ꞵHb+AIR: m=50, n=14, N=10. D) Diagram of the CV of an AP propagating along an axon: distance along the axon (Δx) and the time needed for an AP to pass that distance (t_2_-t_1_). V represents the membrane potential. E) Model of a propagating AP. The change in V(t) is shown at longitudinal increments along the axon (x_2_) and at neighboring increments immediately before (x_1_) and behind (x_3_). The membrane potential at rest (V_R_) changes over time proportionally to the sum of differences between its value and those of the preceding and superseding increments. Upon reaching the threshold potential (V_T_), a spike occurs, and the membrane potential reaches the peak value (VA). F-G) CV is modulated by V_A_ and V_R_. Our computational model predicts declining CV due to reductions in the peak AP (F) and/or hyperpolarization of the resting membrane potential (G). The CV is quantified as a percentage relative to the control.

To better interpret these findings, we constructed a computational model of the CV by employing an approximation of axonal cable theory (see Methods). Our model examined three processes that determine the CV: the resting membrane potential (V_R_), peak membrane potential (V_A_, the amplitude of AP overshoot), and activation threshold potential (V_T_, potential for Na_v_ activation). We considered V_T_ to be robust to various physiological conditions, as this value primarily depends on the Na_v_ channel type and gating kinetics [41]; therefore, we fixed V_T_ at a constant value of −50 mV in our calculations. We varied V_R_ between −90 and −60 mV and V_A_ between 10 and 30 mV, both of which are considered typical physiological ranges. We normalized the results according to a reference state with V_R_ = −75 mV, V_A_ = 30 mV, and V_T_ = −50 mV. The modeling results showed that decreased CV was associated with either hyperpolarized resting membrane potential or decreased peak action potential (AP) amplitude [**Fig. 2F-G**].

### 4.3 AIR adversely affects input timing, which is restored by high D-ꞵHb concentrations

Changes in the CV magnitude and the resulting differences in the input timing would likely desynchronize the hippocampal circuit. Therefore, we investigated whether the latency of FVs changes during stimulation with 20 pulse trains.

In all treatment groups, the early FVs showed ∼-0.2 ms improvement in latency. During the first 3 min, all experimental groups were comparable and exhibited the same pattern of latency changes. Afterward, the AIR and 0.1 mM D-ꞵHb+AIR groups displayed progressively more prominent delays than the Control group, while the Control and 1 mM D-ꞵHb+AIR groups remained comparable. During AIR, we observed the largest delays [**Fig. 3E-F**, **Fig. 3I, L**; at 15 min, stim. 9-11: +2.69 ± 0.84 ms, Control vs. AIR, p=0.0077; 2.64 ± 0.94 ms, AIR vs. 1 mM D-ꞵHb+AIR, p=0.020]. The increased delays during the stimulation suggest that the detrimental effects of AIR are exacerbated over longer periods.

**Figure 3:**
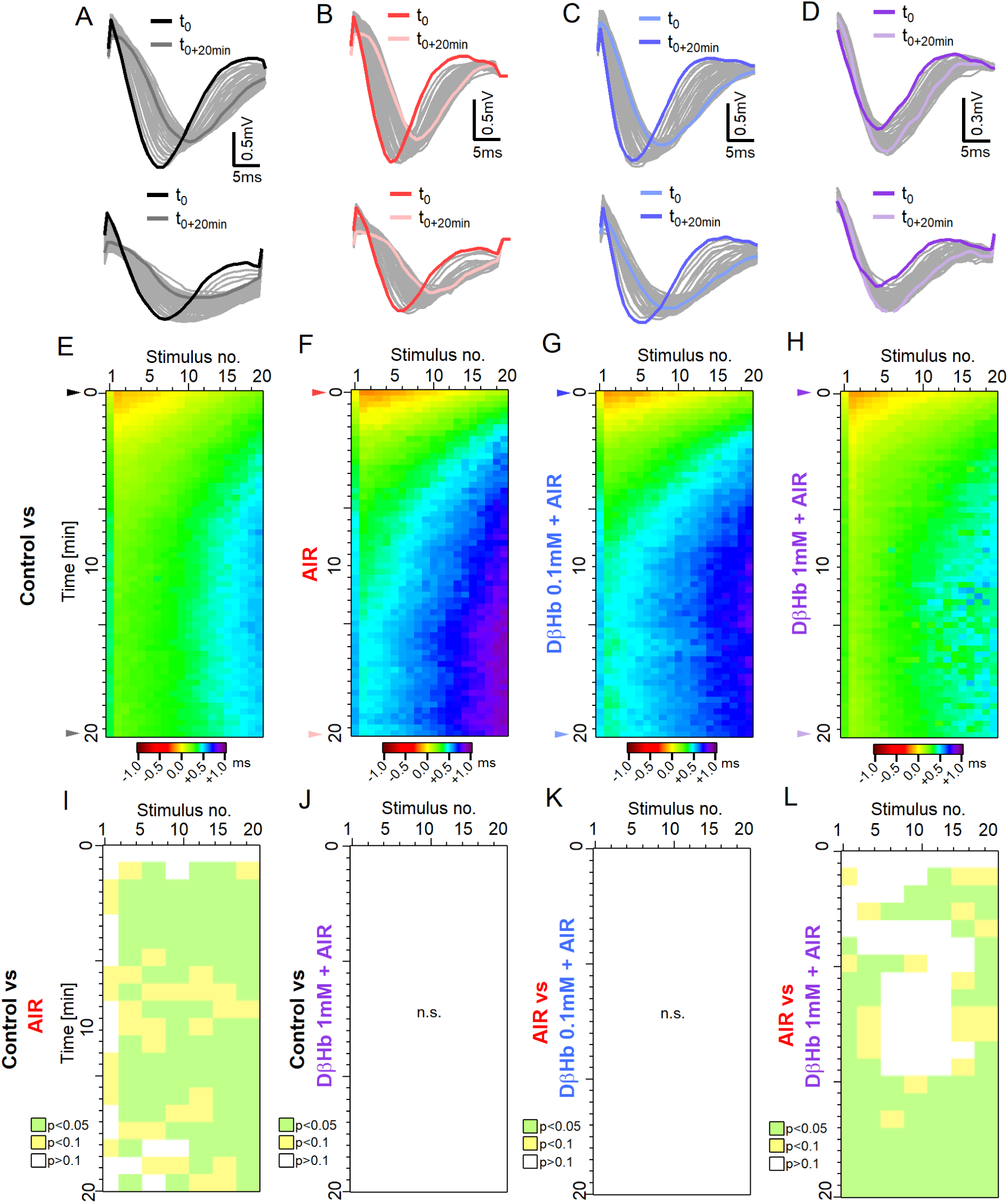
Time delays in axonal firing during train stimulation are increased during AIR and reversed by 1 mM D-ꞵHb. A) Top: Representative example of the FVs recorded at the first stimulus in the control group during stimulation with 20 pulses applied at 25 Hz every 20 s over 20 min. Bottom: FVs recorded in response to the last stimulus in each train. The FVs recorded at t=0 are marked in black; the FVs recorded at t=+20 min are marked in dark gray. B) - D) The same as A) in the AIR, 0.1 mM D-ꞵHb+AIR and 1 mM D-ꞵHb+AIR groups. E) Heatmap of the mean, normalized FV peak latencies recorded in the Control group. Each trial is represented by a new row. The columns represent the time points of successive stimuli. Normalized latencies (0 ms) are color-coded in yellow. Delays (>0 ms) are color-coded in green‒violet, and latency improvements (<0 ms) are color-coded in orange‒brown. n=24, N=22. F) - H) The same as E) for AIR, n=16, N=15; 0.1 mM D-ꞵHb+AIR. n=17, N=15, 1 mM D-ꞵHb+AIR. n=14, N=10. I) Statistical comparison of latencies for the Control (E) and AIR (F) groups, performed based on means of 3 stimuli x 2 trials blocks. p<0.05 in green, p<0.1 in yellow, p≥0.1 in white. J) The same as I) comparing the results of the Control (E) and 1 mM D-ꞵHb+AIR (H) groups. There were no significant differences, p=0.067 to p=0.99. K) The same as I) comparing the results of the AIR (F) and 0.1 mM D-ꞵHb+AIR (G) groups. There were no significant differences, p=0.41 to p=0.99. L) The same as I) comparing the results of the AIR (F) and 1 mM D-ꞵHb+AIR (H) groups.

### 4.4 AIR increases membrane resistance (R_m_) without affecting other intrinsic membrane properties of CA1 pyramidal neurons, and D-ꞵHb treatment during AIR does not rescue R_m_ to normal levels

The above results [**Fig. 2**] indicate that AIR might negatively impact the membrane properties of hippocampal neurons; therefore, we sought to identify the properties most vulnerable to AIR and test D-ꞵHb as a recovery agent. Based on the dose‒response results from the field potential studies, we selected 1 mM D-ꞵHb for use in patch-clamp experiments, testing the effects of AIR and D-ꞵHb on the intrinsic membrane properties of CA1 pyramidal neurons (membrane resistance, R_m_; membrane capacitance, C_m,_ and resting membrane potential, RMP; V_rest_) and their spike thresholds. We found that the R_m_ of the CA1 pyramidal neurons in the AIR group was significantly increased compared with that of the neurons in the control group [**Fig. 4B**, +43.24 ± 3.73%, p=1.41E-07], and this increased value was observed even with the application of 1 mM D-ꞵHb [**Fig. 4B**, +27.86 ± 8.93%, p=0.030], while the results in the AIR and 1 mM D-ꞵHb+AIR groups were similar (p=0.31). Interestingly, neither the C_m_ [**Fig. 4C**, 114.25 ± 9.73, 115.52 ± 9.26, 132.20 ± 9.46, Control, AIR, D-ꞵHb+AIR, respectively, p=0.32], spike threshold [**Fig. 4E**, −46.31 ± 0.76, −44.53 ± 0.80, 44.64 ± 0.73, p=0.17] nor V_rest._ [**Fig. 4F**, −67.26 ± 1.17, −63.57 ± 1.38, −65.11 ± 0.99, p=0.093] were significantly affected by either the AIR or D-ꞵHb+AIR treatments.

**Figure 4:**
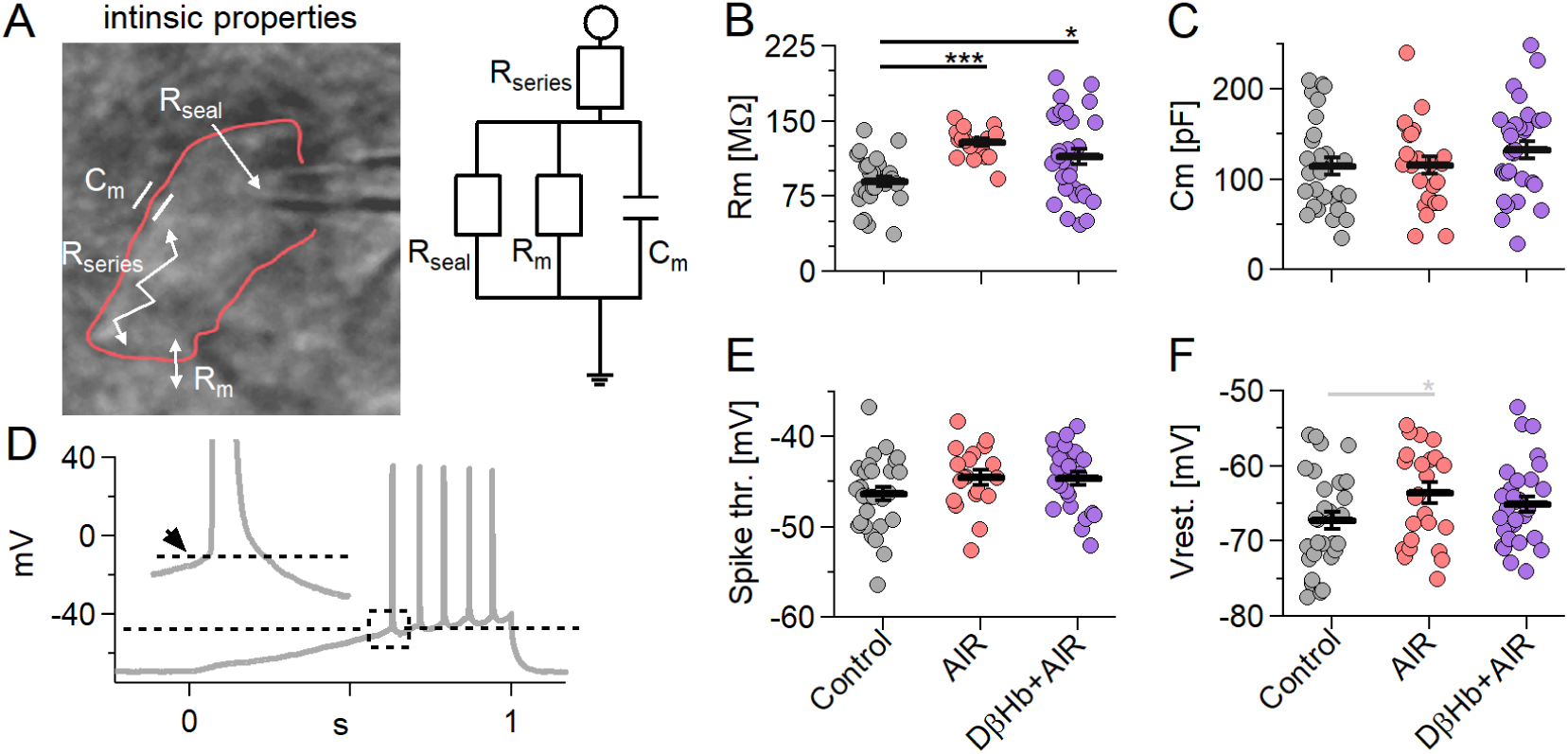
Membrane resistance (R_m_) increases during AIR and is not reversed by D-ꞵHb. Other intrinsic properties did not change under either condition. A) Left: A CA1 pyramidal neuron recorded in whole-cell patch-clamp mode (outlined in red), with resistances and capacitance denoted within the cell. Right: A simple electric circuit diagram of a cell with the same resistances. B) Scatter plots of CA1 pyramidal neuron Rm compared among the experimental groups. Each colored circle represents a value recorded in a single cell. Control in gray, AIR in light red, 1 mM D-ꞵHb+AIR in purple. Vertical black bars represent the mean ± SEM. C) Scatter plots of CA1 pyramidal neurons C_m_ compared among the experimental groups. The plots and coloring are identical to B). There were no significant differences (p=0.32). D) Representative example of a 300 pA ramp current injection performed in a control CA1 pyramidal neuron. The dashed line marks the onset (spike threshold) of the first AP. The inset shows the magnified appreciate, with the arrow pointing to the onset. E) Scatter plots of CA1 pyramidal neuron spike thresholds compared among the experimental groups. The plots are identical to B). There were no significant differences (p=0.17). F) Scatter plots of CA1 pyramidal neurons RMP (V_rest_) compared among the experimental groups. The plots and coloring are identical to B). There were no significant differences (p=0.09). Control n=28, N=22; AIR n=25, N=17; 1 mM D-ꞵHb+AIR; n=30, N=16. In black, *p<0.05; **p<0.01; ***p<0.001; in gray *p<0.1.3.3 AIR adversely affects input timing, which is restored by high D-ꞵHb concentrations.

### 4.5 AIR increases the frequency of spontaneous excitatory postsynaptic currents (sEPSCs) at CA1 synapses and mildly increases sEPSC amplitudes. D-ꞵHb does not reverse the increase in frequency but mildly decreases sEPSC amplitudes

Recovering the membrane back to the resting potential during synaptic transmission is the most energy-consuming neuronal process (∼% of total ATP produced [42]); therefore, we investigated changes in the frequency and magnitude of spontaneous EPSCs that might explain the ∼35% decrease in fEPSP amplitudes reported above [**Figs. 1-3**].

AIR significantly increased the frequency of sEPSCs, with approximately twice as many events observed in the AIR group than in the control group [**Fig. 5A, B**; 0.53 vs. 0.96 Hz, +0.42 ± 0.12 Hz, p=0.0031]. Interestingly, D-ꞵHb failed to rectify the increased sEPSC frequency [**Fig. 5A, B**; D-ꞵHb+AIR 0.93 vs. AIR 0.96 Hz, 0.021 ± 0.15 Hz, p=0.99; D-ꞵHb+AIR vs. Control, +0.40 ± 0.12 Hz, p=0.0091]. However, the amplitude, rise time, and sEPSC decay time did not differ significantly among the groups [**Fig. 5C-E**].

**Figure 5:**
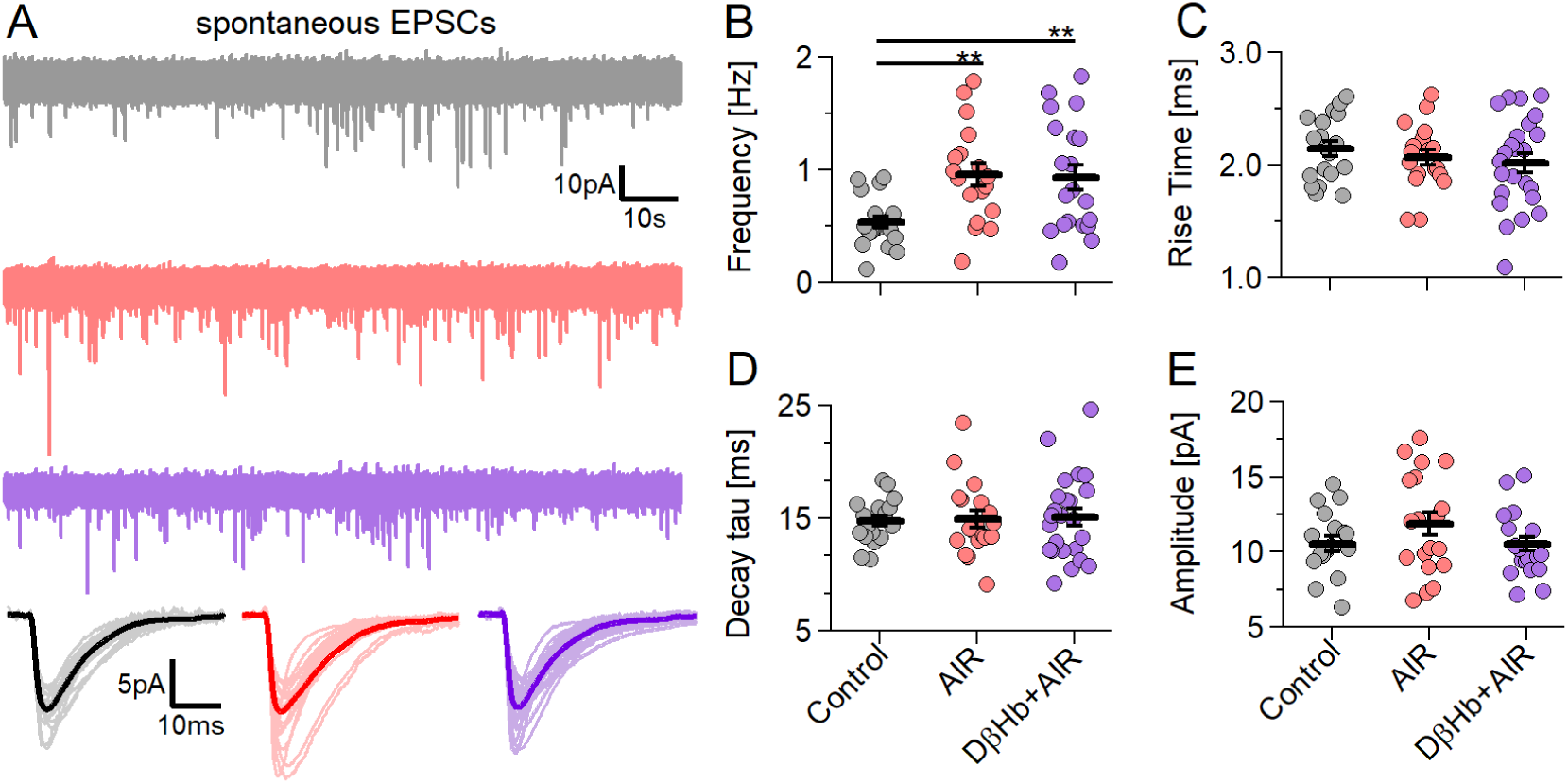
AIR increased sEPSC frequency without altering sEPSC properties, and D-ꞵHb administration did not reverse this effect. A) Representative examples of the first 2 min of sEPSC waveforms recorded in CA1 pyramidal neurons, voltage clamped at V_h_= −70 mV (Control in gray; AIR in red; 1.0 mM D-ꞵHb+AIR in purple). Bottom: averaged sEPSC waveforms from individual cells (light color) and the group mean (full color). B) Scatter plots of the sEPSC frequency recorded in CA1 pyramidal neurons compared among the experimental groups. Each colored circle represents the frequency recorded in a single cell. The labels are the same as in A). Vertical black bars represent the mean ± SEM. *p<0.05. C) Scatter plots of the sEPSC 20-80% rise times, labeled as in B). There were no significant differences (p=0.51). D) Scatter plots of the sEPSC decay tau times, labeled as in B). There were no significant differences (p=0.98). E) Scatter plots of the sEPSC amplitudes, labeled as in B). There were no significant differences (p=0.21). Control n=19, N=19; AIR n=19, N=14; 1 mM D-ꞵHb+AIR, n=23, N=16. **p<0.01.

Given the change in R_m_, the lack of significant changes in the amplitudes is surprising. Several studies utilizing different metabolic states, such as food deprivation [43] and inhibition of glycolysis [44], showed increased miniature EPSC (mEPSC)/sEPSC amplitudes; therefore, we decided to investigate the sEPSCs in more detail. In most central synapses, the distribution of mEPSC/sEPSC amplitudes shows several peaks related to the simultaneous release of multiple quanta of a neurotransmitter [45], with each subsequent peak linked to an increasing number of concurrent sEPSCs. According to the multiplicative nature of this process, we assumed that small changes in amplitudes [**Fig. 5E**] would scale up. We detected 2-4 distinct peaks in ∼85% of the analyzed cells [**Fig. S3A-C**], with no differences in the number of cells with multipeak distributions among the groups [**Fig. S3C**; 89.5% in Control; 84.2% in AIR, 87.5% in D-ꞵHb+AIR]. In line with our initial assumptions, the scaled amplitudes revealed differences among the groups [**Fig. S3E**]: the amplitudes in the AIR group were significantly higher than those in the D-ꞵHb+AIR group [p1: p=0.048; p2: p=0.0054; p3: p=0.0038; p4: p=0.021] and differed from those in the control group at peak 3 [p=0.007]. The Control and D-ꞵHb+AIR groups did not show significant differences at any point [**Fig. S3E]**, despite the elevated sEPSC frequency in the D-ꞵHb+AIR group.

We validated these findings by comparing the normalized amplitude distributions among the different groups [**Fig. S3F-I**]. Within the range correlating to peaks 2–4 (sEPSC of ∼20 pA or greater), significantly more large events were observed in the AIR group (15.19% of total) than in the control (7.65%) or D-ꞵHb+AIR (9.48%) groups [**Fig. S3F-I**], while the results in the control and D-ꞵHb+AIR groups were essentially comparable. This result suggests that during AIR, the elevated frequency of events increased the probability of multi–vesicular events occurring, which was further exacerbated by the ∼20% increase in R_m_. On the other hand, D-ꞵHb+AIR produced a distribution resembling the control, as if the vesicles had lower glutamate content, as suggested by the literature [46,47].

### 4.6 AIR slightly changes the firing pattern of hippocampal pyramidal neurons and causes increased membrane depolarization with input. These effects are reversed by D-ꞵHb, which also increases AP amplitudes

The action potential (AP) generation and subsequent recovery of V_m_ to resting levels require a large part of the energy budget of pyramidal neurons (∼41% of all produced ATP [42]). Therefore, we investigated whether pyramidal neurons retain their input‒output curves (I-O curve) during AIR and how D-ꞵHb alleviates AIR effects.

First, we focused on the firing frequency [**Fig. 6A**]. Cells in the AIR group showed increased firing at lower current injections than those in the D-ꞵHb+AIR group [**Fig. 6B**, 75–150 pA, p=0.0038 to 0.050], but with the same maximal firing rate. These results suggest that the cells activate earlier in the AIR condition than in the control or D-ꞵHb+AIR conditions. To test this further, we measured the maximum depolarization obtained by each neuron during the final 5 ms of each current injection step [**Fig. 6D**]. AIR neurons had slightly higher depolarization with each injection step compared to control neurons [**Fig. 6E, F**; p=0.0027 to 0.048] or D-ꞵHb+AIR [**Fig. 6E, F**; p=0.0029 to 0.042].

**Figure 6:**
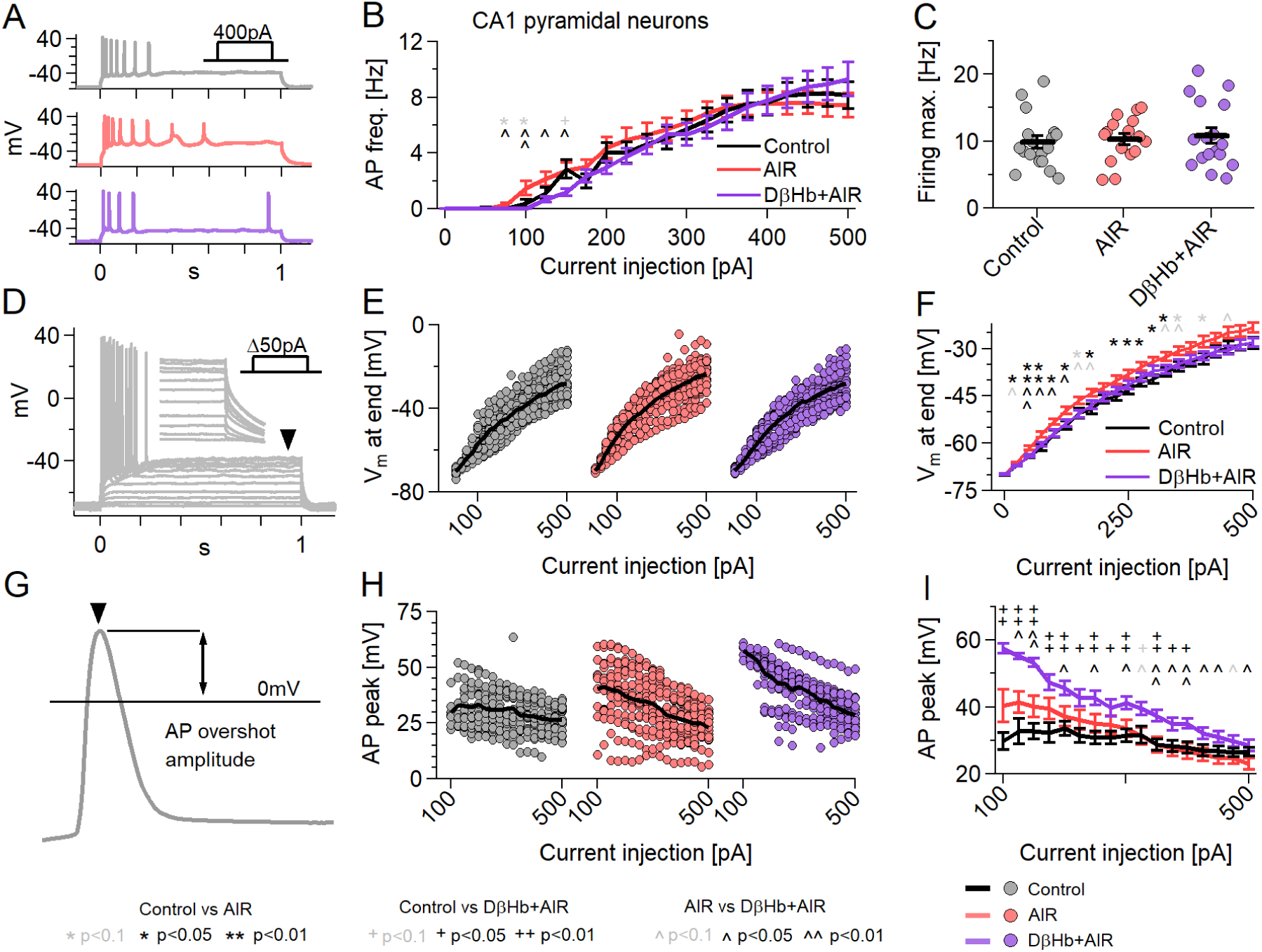
Neuronal firing is mildly affected by AIR and reversed by D-ꞵHb, with D-ꞵHb increasing AP overshoot amplitudes. A) Representative examples of responses to 400 pA square current injections into control (gray), AIR (red), and 1 mM D-ꞵHb+AIR (purple) CA1 pyramidal neurons, held at V_h_=-70 mV. B) I-O curves of CA1 pyramidal neurons receiving 21, Δ+25 pA injections (max. 0 pA; V_h_=-70 mV). Circles represent the group mean ± SEM. * Control vs. AIR, + Control vs. 1 mM D-ꞵHb+AIR, ^ AIR vs. 1 mM D-ꞵHb+AIR. *,+,^ in black p<0.05; **,++,^^ in black p<0.01; *,+,^ in gray p<0.1; ANOVA) Maximum firing rate of the pyramidal neurons in B). Circles represent single neurons; black bars represent group mean ± SEM. There were no significant differences (p=0.778, ANOVA). D) Example of responses to Δ+25 pA square current injections into a control pyramidal neuron. The arrowhead and inset mark the final membrane depolarization (V_m_, last 5 ms of the step). E) Vm at the current injections from B). F) Group mean ± SEM of V_m_ from E). Statistical tests and labels are identical to B). Control n=23, N=20; AIR n=20, N=14; 1 mM D-ꞵHb+AIR n=22, N=16. G) Example of AP overshoot measurement. The black line marks V_m_= 0 mV. H) Averaged AP overshoot amplitudes from B). I) Group mean ± SEM of overshoot amplitudes from H). Statistical tests and labels are identical to B). In all comparisons, Control n=23, N=20; AIR n=20, N=14; 1 mM D-ꞵHb+AIR, n=22, N=16.

Next, we investigated the average AP peak amplitudes [**Fig. 6G**], per step [**Fig. 6H**]. Surprisingly, D-ꞵHb+AIR neurons showed larger amplitudes, which were consistently elevated throughout the injection period, than control or AIR neurons, while AIR neurons remained comparable to control neurons [**Fig. 6I**, p=0.0017 to 0.036, AIR vs. Control; p=0.0093 to 0.044, AIR vs. 1 mM D-ꞵHb+AIR].

### 4.7 AIR increases the AP decay time, while D-ꞵHb accelerates the AP rise time without reversing the AIR-induced slower AP decay

We next proceeded to investigate other AP properties: the decay time (from the overshoot peak to 0 mV) [**Fig. 7A-C**] and rise time (from 0 mV to the overshoot peak) [**Fig. 7D-F**]. We found that control neurons had significantly lower average AP decay times than AIR [**Fig. 7B, C**; p=0.047 to 0.0019] and D-ꞵHb+AIR neurons [**Fig. 7B, C**; p=0.041 to 0.0023] for almost all current injection steps. Moreover, AIR and D-ꞵHb+AIR cells did not significantly differ at any point [**Fig. 7B, C**; p=0.32 to 0.99]. D-ꞵHb+AIR cells demonstrated faster rise times than control cells [**Fig. 7E, F**; p=0.000094 to 0.0025] and AIR cells [**Fig. 6E, F**; p=0.047 to 0.0052] for most injection steps. Interestingly, AIR cells exhibited slightly faster AP rise times exclusively at the lowest current injection times [**Fig. 4N, O**; p=0.012 to 0.035]. These slightly faster APs were likely due to earlier activation of the cells, as shown in [**Fig. 6B-C, E-F**].

**Figure 7:**
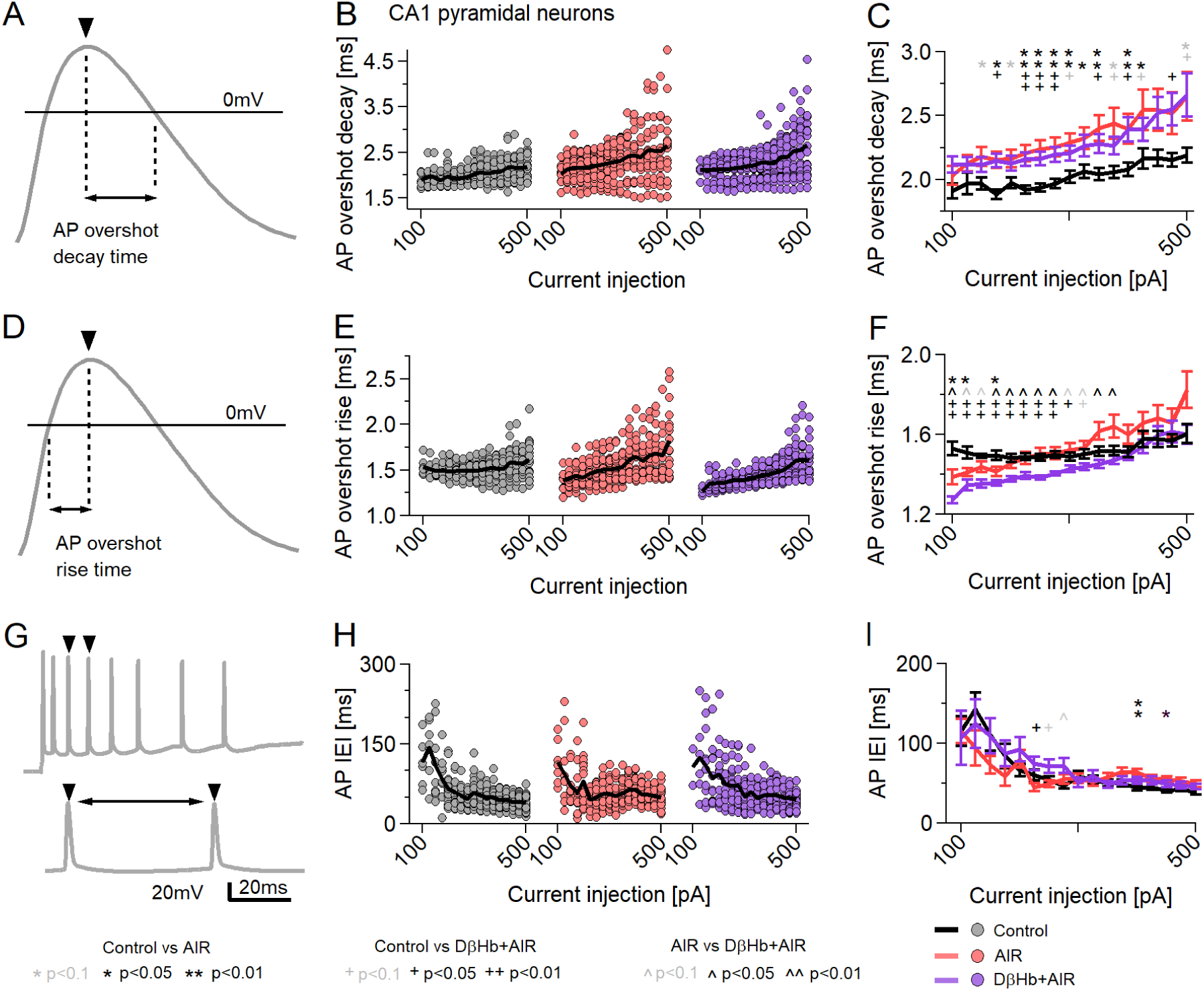
AP decay times change during AIR and D-ꞵHb + AIR conditions, but D-ꞵHb + AIR APs have faster rise times. A) Example of AP decay time measurement. The black line marks V_m_= 0 mV. B) Mean AP decay times for the current injections. Circles represent averaged amplitudes in single neurons. The black lines represent group means. C) Group mean ± SEM of AP decay times from B). Statistical tests and labels are identical to B). D) Example of AP decay time measurement. The black line marks V_m_= 0 mV. E) Averaged AP rise times for the current injections in B). Circles represent averaged amplitudes in single neurons. The black lines represent group means. F) Group mean ± SEM of AP rise times from B). Statistical tests and labels are identical to B). G) An example of the AP IEI measurement. The double arrow marks the latency between AP peaks. H) Averaged IEI recorded during the injections in B). Each point represents the average IEI recorded at each step in a single neuron. The black line represents the group mean. I) Group mean ± SEM of averaged IEI. *,+,^ p<0.05; ANOVA. In all comparisons, Control n=23, N=20; AIR n=20, N=14; 1 mM D-ꞵHb+AIR, n=22, N=16.

Finally, as we did not observe significant differences in the firing frequency among the groups [**Fig. 6B, C**], we examined variations in the timing of AP firing based on the intervals between consecutive APs (AP interevent interval, AP IEI) [**Fig. 7G**], but we did not observe obvious changes in the AP IEI [**Fig. 7H, I].**

### 4.8 AP adaptation is impaired during AIR, with AP amplitudes showing steeper declines, and is not reversed by D-ꞵHb

The AP overshoot amplitudes, decay times, and rise times demonstrated steeper slopes in the AIR and D-ꞵHb+AIR groups than in the control group. The AP properties during neuronal firing undergo dynamic changes [48], commonly referred to as spike adaptation or spike-frequency adaptation [48], and in our experiments, we observed that GLUT4 antagonism intensified these changes.

To test this observation, we compared the peak amplitudes of the first APs [**Fig. S4A-D**], the averaged APs [**Fig. 6G-I**, **Fig. S4E**] and the final APs during current injections [**Fig. S4F**]. The first APs represent neuronal firing after rest (10 s between subsequent injections), while the average and final APs represent firing after energy expenditure. As expected, the first APs displayed minimal changes in amplitude during injections [**Fig. S4D**], with larger amplitudes observed in the D-ꞵHb+AIR group than in the control group. Moreover, although elevated amplitudes were observed in the AIR group, the results did not differ significantly from those in the control or D-ꞵHb+AIR groups. For the final APs, we observed a sharp decline in amplitude, which scaled with the injected current [**Fig. S4F**], in both the AIR and D-ꞵHb+AIR groups. Lower amplitudes were observed in the control group than in the D-ꞵHb+AIR group for 100-300 pA injections (p=0.040 to p=6E-05), and the results remained comparable to those in the AIR group until the final, large, 400-500 pA injections (p=0.032 to p=0.0011), when the APs in the AIR group decreased significantly [**Fig. S4F**]. The results in the AIR and D-ꞵHb+AIR groups remained consistently different. Linear fits to the average AP amplitudes revealed significantly steeper declines in the AIR and D-ꞵHb+AIR groups [**Fig. S4G, H**; p=0.00039, AIR vs. Control; p=0.0012, D-ꞵHb+AIR vs. Control], indicating that prolonged neuronal activity is heavily affected by AIR and D-ꞵHb does not provide direct recovery.

### 4.9 Changes in AP decay and rise times correspond to differences in the widths of FVs

In our initial analysis of the field potential data, we focused on the amplitudes of the biological signals. Consistently, we found no significant changes in the FV amplitudes [**Figs. 1F, 2C, 3E-G**], which supports our findings on AP overshoot amplitudes [**Fig. 6G-I**] (except in the D-ꞵHb+AIR group). Based on these results, we investigated whether the changes in the AP decay and rise times corresponded to changes in the FV widths during train stimulation [**Fig. S5**].

Consistent with our findings on AP decay/rise times [**Fig. 7A-F**], the widest FVs during the stimulation were consistently observed in the AIR group, followed by the control group, with the narrowest FVs observed in the 1 mM D-ꞵHb+AIR group [**Fig. S5D**]. We found differences between the AIR and control groups (p=3.94E-06 to 0.049) and AIR and 1 mM D-ꞵHb+AIR groups (p=4.04E-05 to 0.046) throughout the whole stimulation period, while the results in the control and 1 mM D-ꞵHb+AIR groups were not different (p=0.079 to 0.99), supporting our AP decay/rise time findings [**Fig. 6G-I**].

### 4.10 With abolished GABAergic inhibition, AIR strongly decreased the firing rates of CA1 pyramidal neurons and FSIs, which were not rescued by D-ꞵHb

We next investigated the properties of CA1 pyramidal neurons and FSIs after pharmacologically abolishing AMPA/kainate excitatory (2,3-dioxo-6-nitro-7-sulfamoyl-benzo-quinoxaline; NBQX) and fast GABAergic inhibitory (gabazine) transmission [**Fig. S6**].

We first recorded the firing frequency of CA1 pyramidal neurons [**Fig. S6A, B**] at different activation levels [**Fig. S6C, D**] and found that AIR prominently decreased their firing rate when compared with the control [**Fig. S6C**, p=0.024 to 0.0053]. Surprisingly, the application of 1 mM D-ꞵHb during AIR resulted in a stronger reduction in the firing rate than in the control group [**Fig. S6E**, p=0.000027 to 0.035]. The AIR and D-ꞵHb+AIR results remained similar throughout the injections [**Fig. S6E**]. However, all groups differed significantly with respect to the maximum firing rate, with the highest firing rate observed in the control group (31.03 ± 3.43) and the lowest firing rate observed in the D-ꞵHb+AIR group (13.17 ± 2.05).

Next, we examined CA1 FSIs using the same drug conditions [**Fig. S6E, F**]: AIR decreased the firing rate of FSIs compared with the control [**Fig. S6G**, p=0.012 to 0.048], which was not reversed by D-ꞵHb [**Fig. S6H**, p=0.00017 to 0.035]. For pyramidal neurons, the FSIs in the AIR and D-ꞵHb+AIR groups remained mostly comparable [**Fig. S6G**]. None of the groups differed in the maximum firing rate [**Fig. S6H**, p=0.068].

### 4.11 The Hodgkin-Huxley model predicts that impairments in Na^+^/K^+^ ATPase activity result in more depolarized V_rest_, lower AP overshoot (V_a_), and increased neuronal firing

Here, we aim to explain the previous observations of dysregulated firing dynamics in response to AIR. We hypothesize that indinavir restricts the energy supply of neurons and thus impairs the neuronal Na^+^/K^+^ ATPase. To test this hypothesis, we developed a computational model of an isolated CA1 neuron in the absence of inhibition and toggled Na^+^/K^+^ ATPase activity through the enzyme kinetic parameter v_ATPase_^max^ (see Supplementary Methods, **Fig. S7**). **Fig. S7A, B** (at two different temporal scales) shows the predicted changes in neuronal firing dynamics when the Na^+^/K^+^ ATPase is impaired. This impairment predicted the depolarization of the resting membrane potential [**Fig. S7C**] and is consistent with the trends of our I/O curves [**Fig. 4F**]. Furthermore, in this depolarization model, the magnitude of the membrane potential shift needed for an AP is reduced. Consequently, the model predicted increased firing frequency [**Fig. S7D**]. Finally, the model predicted that the Na^+^/K^+^ ATPase should act more slowly to recover ion gradients after an AP, causing the amplitudes of the subsequent APs to decline progressively with time [**Fig. S7E, F**]. This amplitude decrease is consistent with the trends in the measured CV [**Fig. 2**].

## 5. Discussion

Central insulin resistance impairs cognitive performance and promotes hippocampal neurodegeneration and memory loss [49,50]. Our present results highlight the adverse effects of GLUT4 inhibition on synaptic function and LTP, in accord with findings that acute and chronic blockade of GLUT4 are detrimental to hippocampus-mediated memory tasks, thus suggesting that GLUT4 is critical for the acquisition and consolidation of memory [8,14]. Memory formation promotes GLUT4 incorporation into the membrane, resulting in increased glucose flux into neurons [14,51]. This process enables increased metabolic support during high-demand activities. Crucially, most GLUT4s localize in the perikaryon [52,53], likely near axosomatic synapses, which are associated with synaptic plasticity and long-term memory (LTM). Indeed, prolonged blockade of GLUT4 glucose transporters in the brain results in impaired formation of LTM [14] and decreased levels of hippocampal brain–derived neurotrophic factor (BDNF) [54]. Moreover, hypoglycemia negatively affects memory performance and cognition [55,56; however, see 14].

In this study, we found that AIR increased the frequency of sEPSC events, a finding which is consistent with previously published data showing similar increases during food deprivation [44], inhibition of glycolysis [43] and ischemia [57], likely due to hypoglycemia-driven Ca^2+^ accumulation at presynaptic terminals [43,57]. These studies also suggest that presynaptic glycolysis is essential for maintaining synaptic transmission, even at low frequencies, and that mitochondrial respiration cannot compensate for these effects [43,57]. Normally, glycolysis requires approximately one-third of the energy used at presynaptic terminals, thus sustaining low-frequency transmission. In addition, impaired glycolysis leads to slower, broader, and smaller AP waveforms and depolarization of the resting membrane potential [44], which is consistent with our results. Interestingly, D-ꞵHb+AIR treatment, which showed the same increase in the frequency of sEPSC events as AIR, led to the lowest sEPSC amplitudes [**Fig. S3E**]. This result suggests a reduction in the number of synaptic vesicles and is consistent with reports of reduced conversion of glutamate to aspartate during ketosis [46,47].

Although the results of our computational models are consistent with the observed changes in the resting-state membrane potential and firing frequency during AIR, there were seemingly contradicting results with respect to the AP peak amplitude. Specifically, we found no significant changes in the average AP overshoot amplitudes during AIR, and our model predicted a decrease due to energy constraints. In the model, we did not consider a progressive decline in available energy, which is suggested by the steeper AP adaptation [**Fig. S4**]. Moreover, neurons may compensate for such energetic limitations by increasing R_m_ (as reported in 44), while our model assumes that R_m_ remains unchanged. When those factors are accounted for, the model and experimental data agree: during periods of extensive activity, energy reserves in neurons are progressively depleted and, provided that the neuron rapidly fires multiple APs, the negative effects of hypoglycemia overcome compensatory R_m_ increases; then, the AP amplitudes decrease below control levels [**Fig. S4**]. Furthermore, we used the maximal Na^+^/K^+^ ATPase activity to represent ATP availability and considered literature estimates of kinetic and thermodynamic rate constants. A more direct approach might consider ATP concentration as an input variable, in cases in which such experimental data are available.

Our model aims to establish the underlying causes of the changes in CV, and we investigated V_rest_ and AP peak amplitude as two critical factors. The model explains CV recovery and CV increases during D-ꞵHb+AIR treatment, as the measured AP peak amplitudes remain consistently elevated and V_rest_ changes are negligible, but fails in the case of AIR alone, with the AP amplitudes remaining comparable to the control levels and V_rest_ tending to depolarize. This discrepancy might reflect the simplistic nature of our model, which neglects complex ion dynamics, altered membrane permeability and synaptic processes.

The ketogenic diet has been used to treat epilepsy for decades and has been considered in many therapeutic applications [32]. Possible mechanisms include modulation of ATP-sensitive K^+^ channels (K_ATP_), free fatty acid receptor 3 (FFAR3/GPR41) activation, promotion of GABA synthesis, and epigenetic modifications [58]. The action of D-ꞵHb in modulating potassium flux through K_ATP_ channels may be most relevant in the present context [59] since K_ATP_ activation results in decreased neuronal firing rates [60,61] and prevents picrotoxin-induced epileptiform activity in the hippocampus [62]; thus, this action may explain the present findings that firing rates and membrane depolarization remained at control levels in the presence of 1 mM D-ꞵHb during AIR (despite elevated R_m_).

D-ꞵHb is a precursor of glutamine, which promotes increased GABA levels, as seen in studies on the ketogenic diet in patients with epilepsy [63]. D-ꞵHb also directs glutamate toward GABA production by reducing the conversion of glutamate to aspartate [46,47]. In addition, D-ꞵHb directly affects the vesicular glutamate transporter 2 (VGLUT2) by inhibiting Cl^−^ dependent glutamate uptake. VGLUT2 is critical for glutamate excitatory transmission in CA3-CA1 [64], and conditional VGLUT2 knockout results in impaired spatial learning and reduced LTP [64]. Together with the increased formation of GABA and/or GABA derivatives, VGLUT2 inhibition could reduce glutamate excitatory transmission [65].

The present results have important implications for the treatment of conditions such as type 2 diabetes, metabolic syndrome, and seizure disorders that involve insulin hyposensitivity. According to NIH estimates, ∼38% of Americans display symptoms of prediabetes [66], and this percentage is expected to increase. With longer average lifespans and demographic changes leading to an increase in the aging population, obtaining insights into the outcomes of metabolic dysfunction and its potential treatment has become a crucial issue. Our results provide unique insight into the potential use of KBs as an inexpensive and relatively risk-free treatment for metabolic disorders and insulin resistance. Medical researchers can use this information to discover new targets for treatment and develop more effective therapies. Further research in this area could inform novel treatment approaches that address the complex effects of metabolic disorders on brain health.

## 6. Materials and Methods

### 6.0 Mice

Breeding pairs of transgenic GAD2Cre/GCamp5gTdTomato, Thy1/GCamp6fTdTomato, and C57BL/6J mice were originally obtained from The Jackson Laboratory (stocks 010802, 024477, 024339) or Charles River Laboratories and bred in-house under standard conditions of a 12-12 hour light-dark cycle, with food and water available *ad libitum*. The University of Rochester School of Medicine and Dentistry Institutional Animal Care and Use Committee approved all experiments. Mice of both sexes aged between 30 and 60 days (P30-P60) were used in all experiments.

### 6.1 Electrophysiology – slice preparation

Coronal brain slices containing the hippocampus were used in all recordings. Mice were deeply anesthetized with a mixture of isoflurane and air (3% v/v) and decapitated, and their brains were extracted and cut using a Leica VT1200S vibratome into 300 μm-thick slices in ice-cold ACSF solution containing (in mM): 230 sucrose, 1 KCl, 0.5 CaCl_2_, 10 MgSO_4_, NaHCO_3_, 1.25 NaH_2_PO_4_, 0.04 Na-ascorbate, and 10 glucose; 310 ± 5 mOsm, pH adjusted to 7.35 ± 5 with HCl, and gassed with carbogen (95% O_2_, 5% CO_2_) for at least 30 min before use. The slices were then transferred to a BSC-PC submerged slice chamber (Harvard Apparatus, USA) filled with artificial cerebrospinal fluid (ACSF) containing (in mM): 124 NaCl, 2.5 KCl, 2 CaCl_2_, 2 MgSO_4_, 26 NaHCO_3_, 1.25 NaH_2_PO_4_, 0.04 Na-ascorbate, and 10 glucose; 305 ± 5 mOsm/kg, pH adjusted to 7.35 ± 0.05 mOsm/kg, gassed with carbogen and prewarmed to 32 °C. The slices were allowed to recover in warm ACSF for 15 min; then, the heating was switched off, and the chamber was gradually cooled to room temperature (RT; 23 ± 2 °C) over 45 min.

Following recovery, individual slices were transferred to another BSC-PC chamber filled with gassed ACSF and a mixture of drugs (indinavir 0.1 mM; indinavir 0.1 mM and either 0.1- or 1-mM D-ꞵHb), where they were incubated for 60 ± 15 min. Next, the slices were transferred to a submerged recording chamber mounted on an upright microscope stage (E600FN, Nikon, Japan) equipped with infrared differential interference contrast (IR-DIC) filters. The slices were kept at RT and superfused continuously with gassed ACSF delivered by a peristaltic pump at a steady rate of ∼2 ml/min.

### 6.2 Field potential recordings

Evoked field potential responses were triggered with an isolated pulse stimulator (Model 2100, AM Systems, USA) using a CBARC75 concentric bipolar electrode (FHC, USA) placed between CA3 and CA1. The recording electrode was placed 300 ± μm away from the stimulation electrode for the CA1 FV and fEPSP recordings. During the axonal conduction velocity (CV) recordings, the electrodes were spaced 300 to 900 ± 50 μm apart, and up to 6 different sites were recorded. All stimulation paradigms were applied every 20 s as biphasic rectangular pulses with a total duration of 240 μs, 50 ± 5 V strength (∼60 ± 10% of the maximum response size), and a between-stimuli frequency of 25 Hz (40 ms).

During non-CV recordings, three different types of stimulation were applied: paired, 20 stimuli, and stimuli trains, repeated every 20 s for a total period of 20 min in each case. After each train stimulation, the slice was allowed to recover for 20 min under paired stimulation conditions.

Recording pipettes were pulled from borosilicate glass capillaries (ID 0.86, OD 1.5, BF1-86-10, Sutter Instruments, USA) with a vertical PC-100 puller (Narishige, Japan) and filled with 6% NaCl (in H_2_O, 1.02 M). Only pipettes with a resistance of 0.2-1.0 MΩ were used. Recording pipettes were allowed to rest at the recording site for 10 min to equilibrate before the start of a recording. All recorded signals were acquired using a MultiClamp 700B amplifier (Molecular Devices, USA), Bessel-filtered at 2 kHz, and digitized at a sampling frequency of 10 kHz using an Axon Digidata 1440A digitizer (Molecular Devices, USA).

### 6.3 Whole-cell patch-clamp recordings

Pyramidal neurons in the hippocampal CA1 region were selected for recordings based on their general morphology and position within the pyramidal layer. As an additional verification, recorded cells were screened for glutamate decarboxylase 2 (GAD2) ^+^-driven tdTomato fluorescence. Cells that displayed any level of tdTomato fluorescence were rejected.

Patch pipettes were pulled from borosilicate glass capillaries (BF1-86-10, Sutter Instruments, USA) with a vertical puller (PC100, Narishige, Japan). Pipettes had a resistance of 6–8 MΩ when filled with an internal solution containing (in mM): 136 K-gluconate, 4 disodium adenosine 5′-triphosphate (Na_2_ATP), 2 MgCl_2_, 0.2 ethylene glycol-bis(β-aminoethyl ether)-N,N,N′,N′-tetraacetic acid (EGTA), 10 4-(2-hydroxyethyl)-1-piperazineethanesulfonic acid (HEPES), 4 KCl, 7 di(tris) creatine phosphate, 0.3 trisodium guanosine 5’-triphosphate (Na_3_GTP); 280–290 ± 5 mOsm/kg, titrated to pH 7.3 ± 0.05 with KOH.

During the voltage clamp (VC) recordings, cells were clamped at a holding potential (V_h_) of −70 mV with a Multiclamp 700B amplifier (Molecular Devices, USA). The liquid junction potential was not corrected. The pipette capacitance was compensated for in all recordings. During VC experiments, the series resistance (R_s_) was not compensated. During the current clamp (CC) recordings, a holding current was applied to the cells to maintain the membrane potential at V_h_ = −70 ± 3 mV. All cells were bridge-balanced at 50 % of R_s_. The cell capacitance was not compensated for in either mode. Immediately after establishing the whole-cell configuration, the cell was kept in I=0 mode to monitor V_rest_ over a period of ∼120 to 180 s. The average value of V_rest_ over this period is reported as the neuron’s V_rest_.

During the recordings, every 5 to 10 min, a series of 10 square voltage steps of −10 mV was applied to monitor changes in R_s_ (holding voltage; V_h_ = −70 mV). All whole-cell currents in response to voltage steps were low-pass filtered at 10 kHz with a Bessel filter and sampled at 20 kHz frequency (Axon Digidata 15B, Molecular Devices, USA). For inclusion in the final dataset, recordings had to meet 3 criteria: 1) The R_s_ value after any recording protocol could not be more than 20% larger than the R_s_ value before the protocol; 2) the R_s_ value during any of the recording protocols could not exceed 30 MΩ; and 3) the offset drift by the end of the recordings could not be higher than ± 5 mV.

sEPSCs were recorded in VC at V_h_ = −70 mV, in a single, continuous sweep for 10 to 15 min, low-pass filtered at 2 kHz with a Bessel filter and sampled at 10 kHz frequency.

To establish the spiking threshold, each neuron was current-clamped at V_h_ ∼ −70 ± 3 mV and subjected to a ramp-shaped 200-500 pA current injection with a 1 s duration. The spike threshold was determined as the membrane potential at which the first fully developed AP was visible. Each ramp injection was repeated 10x, and the averaged threshold value is reported for the neuron.

To establish the firing properties of the recorded neurons, each neuron was current-clamped as described previously and subjected to a series of square current injections with a change of 25 pA per step to a maximum of 0 pA, with each step having a 1 s duration and an interstep interval (ISI) of 10 s. If necessary, the holding current was corrected during the protocol to maintain V_h_ at ∼ −70 ± 3 mV. During analyses, only fully formed APs that crossed the 0 mV threshold were included.

All data acquisition was performed using pCLAMP software (Molecular Devices, USA).

### 6.4 Field potential recordings – analyses

During field potential recordings, the biologically relevant signals are superimposed onto the decaying stimulation artifact. To isolate the stimulation artifacts for subtraction from the signal at the end of each recording, we applied 1 μM tetrodotoxin (TTX) to the slice and recorded 30 repetitions of each stimulation paradigm in the presence of TTX.

Recorded traces were analyzed in Igor Pro (Wavemetrics, USA). First, an average of the traces recorded with TTX was subtracted from each trace. The traces were then filtered using a rolling five-point average to remove noise, and the baseline was adjusted to the 1 s period before the stimuli. The amplitudes and latencies of the FVs and fEPSPs were measured at their peaks. Amplitudes were measured against 0 mV baselines, while latencies were measured against the onset of the stimulation. During high-frequency stimulation, cells often do not have sufficient time to fully recover their membrane potentials before the next stimulus is applied, leading to a gradual shift in the baseline; therefore, as a correction, we subtracted the median voltage at +35 to +40 ms after each stimulus from each FV or fEPSP amplitude. FV widths were measured at the peaks of positive deflections in the waveform, with the first peak after the artifact subtraction and the second peak directly before the fEPSP.

All traces were screened for occasional electrical interference, and the affected data points were manually removed from the analysis.

To measure the axonal conduction velocity (CV), we calculated the time shift (Δt, ms) between the stimulus start and the time of the FV peak. Next, the linear distance between the recording site and the stimulation electrode was divided by the measured Δt. We recorded CV values at 3-6 sites over a span of 300-900 μm, with ∼1 μm steps between sites. Each CV recording site served as a replicate for statistical analysis (see Statistics). During CV recording, the stimulation site remained stationary, and the sites were recorded in randomized order.

To calculate changes in the FV peak latency during train stimulations, the median Δt for stimuli 1-3 and traces 1-2 (Δt_N_) was subtracted from each Δt (normalization).

### 6.5 Whole-cell patch-clamp – sEPSC analyses

sEPSC detection was automatized using a convolution-based algorithm in Fbrain 3.03 [67], a customized program in Igor Pro 6 (WaveMetrics, Lake Oswego, USA), kindly provided by the Peter Jonas Lab (IST, Klosterneuburg, Austria). Recorded traces were smoothed by subtracting a 20-term polynomial fitted to the trace and digital notch (60 ± 0.5 Hz) filtered in FBrain before the analysis. The convolved trace was passed through a digital bandpass filter at 0.001 to 200 Hz. The event detection template had a rise-time constant τ = 1.5 ms, a decay τ = 6 ms, and an amplitude of −10 pA. The event detection threshold (θ) was set to 4.2 times the standard deviation of a Gaussian function fitted to the all-point histogram of the convolved trace. All events detected by the algorithm were inspected visually. Events that clearly did not show the fast rise times and exponential decay kinetics of a typical hippocampal EPSC were manually removed. Only cells with a rejection ratio lower than 20% were included in the analysis. The analysis was performed using custom-written macros provided by Dr. Maria Kukley (Achucarro Basque Center for Neuroscience, Bilbao, Spain).

Afterwards, individual sEPSC waveforms were extracted from the trace. Each waveform consisted of a 5 ms segment of the trace before the onset of the sEPSC and a 50 ms segment after the sEPSC onset to capture the full decay of the waveform. Before averaging, all sEPSCs were baseline-adjusted to the pre-onset time by 5 ms. The peak amplitude, 20-80% onset-to-peak rise time, and decay constant τ were measured for the averaged sEPSCs. The decay constant τ was calculated from the monoexponential fit from the sEPSC peak to the final ms of the averaged waveform.

To avoid selection bias during sEPSC extraction, we repeated the Fbrain analysis at least twice for each cell and averaged the results.

### 6.6 Statistics

All data were acquired in randomized sequences, with a maximum of 2 slices (field potential recordings) or 2 cells (patch-clamp recordings) from a single animal per experimental condition to avoid pseudoreplication. All experimental groups had comparable distributions of ages (close to P50 ± 6) and sex. The order of slice preincubation with drugs was randomized for each animal. The numbers of slices or cells and animals used in each experiment are indicated in the figure legends.

Statistical analysis was performed using GraphPad Prism 9.5.1 (GraphPad Software, USA). Significant outliers were removed with the Prism ROUT method at Q = 5%, and the normality of residuals and homoscedasticity were tested for each comparison. If the datasets had normal residuals and equal variances, we used ordinary one-way ANOVA with *post hoc* Holm-Šídák’s test. If the datasets had normal residuals but unequal variances, we used Brown-Forsythe ANOVA with *post hoc* Dunnett’s T3 test. If the datasets were not normally distributed but had equal variances, we used Kruskal-Wallis ANOVA with *post hoc* Dunn’s test. In rare cases where the data were neither normally distributed nor had equal variance, we applied Brown-Forsythe ANOVA with *post hoc* Dunnett’s T3 test. If the dataset consisted of multiple replicates, we used nested one-way ANOVA with *post hoc* Holm-Šídák’s test. The tested groups and p values are indicated in the text or the figure legends. Individual cells or slices are labeled n, replicates are labeled m, and the numbers of animals used are labeled N. If the groups tested are different, we report the p values of *post hoc* tests but omit the p value of the omnibus test. If the groups did not test as different, only the p value of the omnibus test is reported. For scatter plots, each point represents an individual data point (cell, slice, or replicate), and the horizontal bars represent the group mean ± SEM. In all other graphs, we present the mean ± SEM. The heatmaps represent the group means.

## Supporting information

Manuscript

## 7. Acknowledgments

We thank Dr. Vittorio Gallo (Children’s National Research Institute, USA) for sharing his electrophysiology setup, without which this work would not have been possible; Dr. Kieran Clarke (University of Oxford) for providing D-ꞵHb ester and advice; Dr. Joseph Abbah (Children’s National Research Institute, USA) for help during the preliminary stage of the research; Dr. Richard L. Veech and Dr. M. Todd King (Lab of Metabolic Control, NIH/NIAAA, USA) for providing D-ꞵHb ester and advice; Dr. Ting-Jiun Chen (Mt. Sinai Medical Center, USA) for help and advice with setting up the patch-clamp recordings; Dr. Peter Jonas (Institute of Science and Technology Austria, Austria) for providing the FBrain software; Dr Fernando Fernandez (Boston University, USA) for comments on the first draft of the manuscript; Dr Paul Hruz (Washington University School of Medicine in St. Louis, USA) for help with establishing the GluT4 inhibition model; Dr Nandkishore Prakash and Dave Saxon (Children’s National Research Institute, USA) for advice and discussions; Dr Heidi Matos Galicia (National Institute of Neurological Disorders and Stroke, USA) for advice; Dr. Paul Cumming (Bern University Hospital) for critical reading of the manuscript; and members of the Mujica-Parodi laboratory (LCNeuro, USA) for discussions; C.W. also acknowledges support from the Marie-Josée Kravis Fellowship in Quantitative Biology.

This manuscript was posted on a preprint: https://www.biorxiv.org/content/10.1101/2023.08.23.554428v3

## 9. Contributions

**Conceptualization**: BK, BA, CW, LMP, NAS

**Methodology**: BK, BA, FG, AB, JMH, CW, MK, LMP, NAS

**Software**: FG, AB, JMH, BA, CW, MK

**Validation**: BK, BA, CW, LMP, NAS

**Formal Analysis**: BK, BA, CW, FG, AB, JMH

**Investigation**: BK, BA, CW, FG, AB, JMH, LMP, NAS

**Writing – Original Draft**: BK

**Writing – Review & Editing**: BK, BA, CW, FG, AB, VV, JMH, MK, LMP, NAS

**Visualization**: BK, BA

**Supervision**: LMP, NAS

**Funding Acquisition**: LMP, NAS

## 11. Data and materials availability

All Data and Data tables with all analyses and tables with statistics are available at Dryad: https://datadryad.org/stash/share/XnKk8GOJq2V3Frm_KJ3BGkg7rf6P9RZRnnQlW4X5Qso. All code is available on GitHub by following the links provided in the text.

