## Supplementary material for "*D*-*ꞵ*-hydroxybutyrate stabilizes hippocampal CA3-CA1 circuit during acute insulin resistance": Manuscript

Bartosz Kula *et al.*

**This PDF file includes:**

Figs. S1 to S8

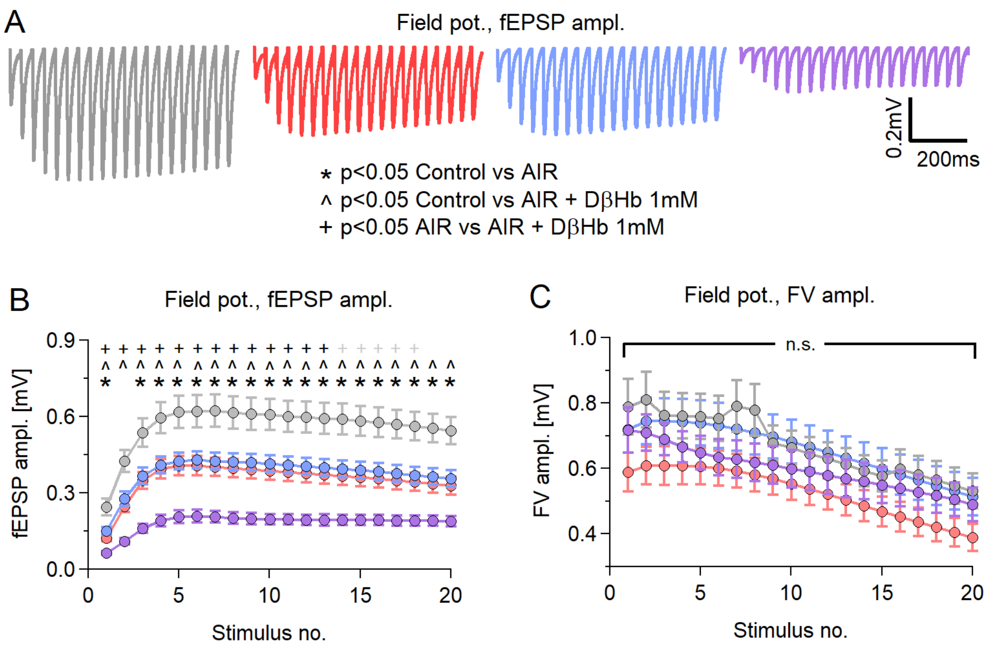


**Figure S1: AIR and D-βHb have the same effects during train and paired stimulation.**

A) Representative examples of averaged fEPSP responses, recorded in each experimental group, to Schaffer collateral stimulation with 20 pulses applied at 25 Hz every 20 s for 20 min, with 280-350 µm distances between electrodes. Control in gray; AIR in red; 0.1 mM D-βHb+AIR in blue; 1 mM D-βHb+AIR in purple. Stimulation artifacts and FVs are removed for clarity. B) Group mean fEPSP amplitudes ± SEM evoked by stimulation, as in A), plotted against the stimulus number. *, +, and ^ indicate statistically significant differences. *<0.05 Control vs. 0.1 mM AIR; #p<0.05 Control vs. 0.1 mM D-βHb+AIR; +p<0.05 Control vs. 1 mM D-βHb+AIR; ^p<0.05 0.1 mM D-βHb+AIR vs. 1 mM D-βHb+AIR; ANOVA. Control, n=24, N=22; AIR, n=16, N=15; 0.1 mM D-βHb+AIR, n=17, N=15; 1 mM D-βHb+AIR, n=14, N=10. C) Group mean FV amplitudes ± SEM recorded during A), plotted against the stimulus number. The labels n and N are identical to (B). There were no significant differences (p=0.091 to 0.34 during the trains, ANOVA).

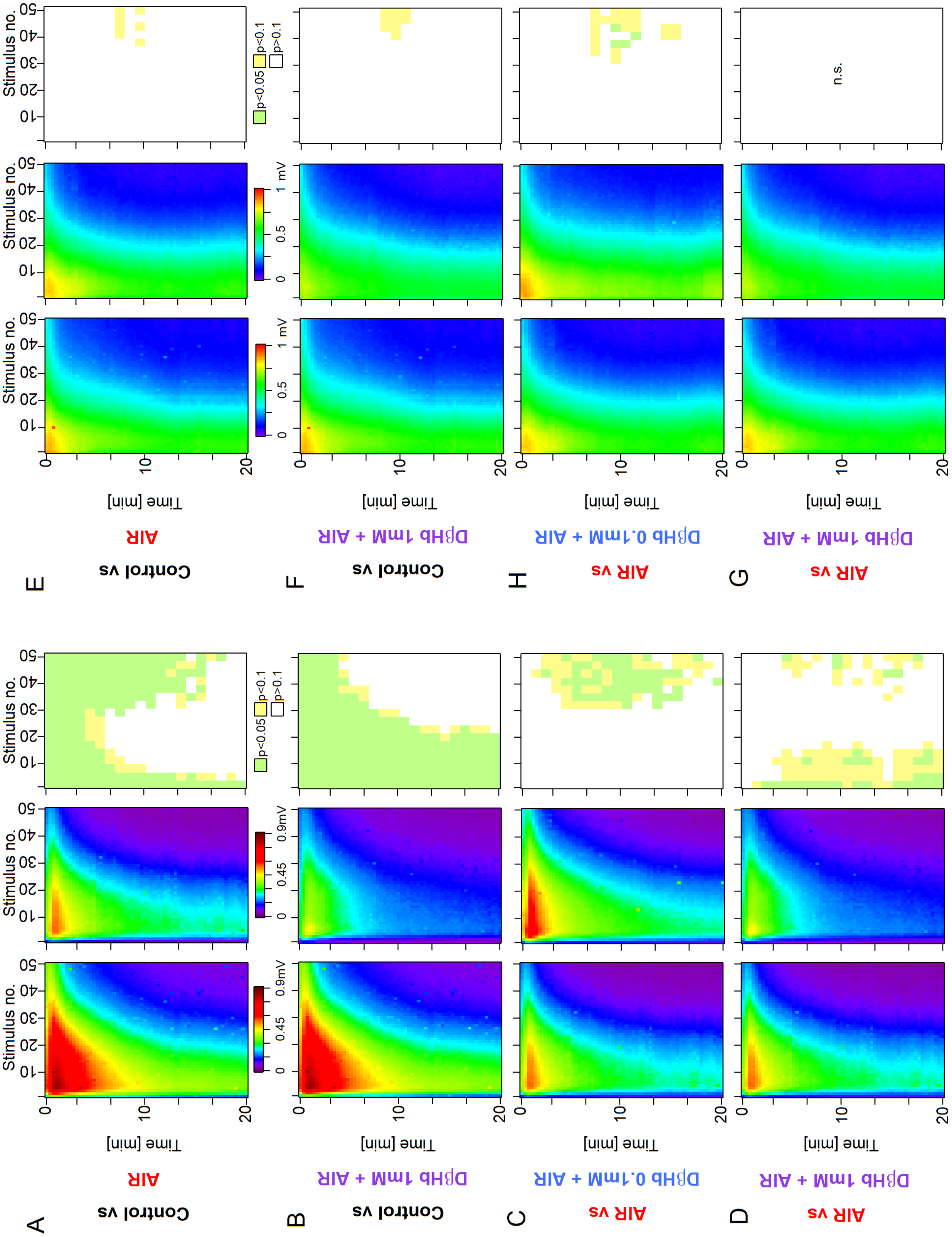
A
Control vs
AIR
B
Control vs
DβHb 1mM + AIR
C
AIR vs
DβHb 0.1mM + AIR
D
AIR vs
DβHb 1mM + AIR
E
Control vs
AIR
F
Control vs
DβHb 1mM + AIR
H
AIR vs
DβHb 0.1mM + AIR
G
AIR vs
DβHb 1mM + AIR
Time [min]
Stimulus no.
0.9mV
0.45
1 mV
0.5
p<0.05
p<0.1
p>0.1
n.s.

**Figure S2: During overtaxing, nonphysiological stimulation, AIR and D-βHb show comparable effects with physiological stimulation paradigms.**

**A)** Heatmaps of mean fEPSP amplitudes recorded over 60 min in the Control and AIR groups and the significance of the differences. The y-axis represents the time points of the trains of 50 stimuli applied at 25 Hz every 20 s, with 280-350 µm distances between electrodes. The x-axis represents the time points of responses to individual stimuli. fEPSP amplitudes are color-coded as a visible light spectrum in the range of 0.1 mV (brown) and 0 mV (violet). Control, n=24 slices, N=24 mice; AIR, n=21, N=18. The groups were compared as 3 stimuli x 3 time blocks, and the p values are summarized on the right-most graph. **B)** The same as A) for the Control and 1 mM D-βHb+AIR groups. Control n=24, N=24; 1 mM D-βHb+AIR n=12, N=10. **C)** The same as A) for the AIR and 0.1 mM D-βHb+AIR groups. AIR n=21, N=18; 0.1 mM D-βHb+AIR; n=18, N=15. **D)** The same as A) for the AIR and 1 mM D-βHb+AIR groups. AIR n=21, N=18; 1 mM D-βHb+AIR n=12, N=10. **E)** Comparison of the mean FV amplitudes between the Control and AIR groups, as in A). The layout, color-coding, and n and N numbers are identical to A). **F)** The same as E) for the Control and 1 mM D-βHb+AIR groups. The layout, color coding, and n and N numbers are identical to B). **G)** The same as E) for the AIR and 0.1 mM D-βHb+AIR groups. The layout, color coding, and n and N numbers are identical to C). **H)** The same as E) for the AIR and 1 mM D-βHb+AIR groups. The layout, color coding, and n and N numbers are identical to D).

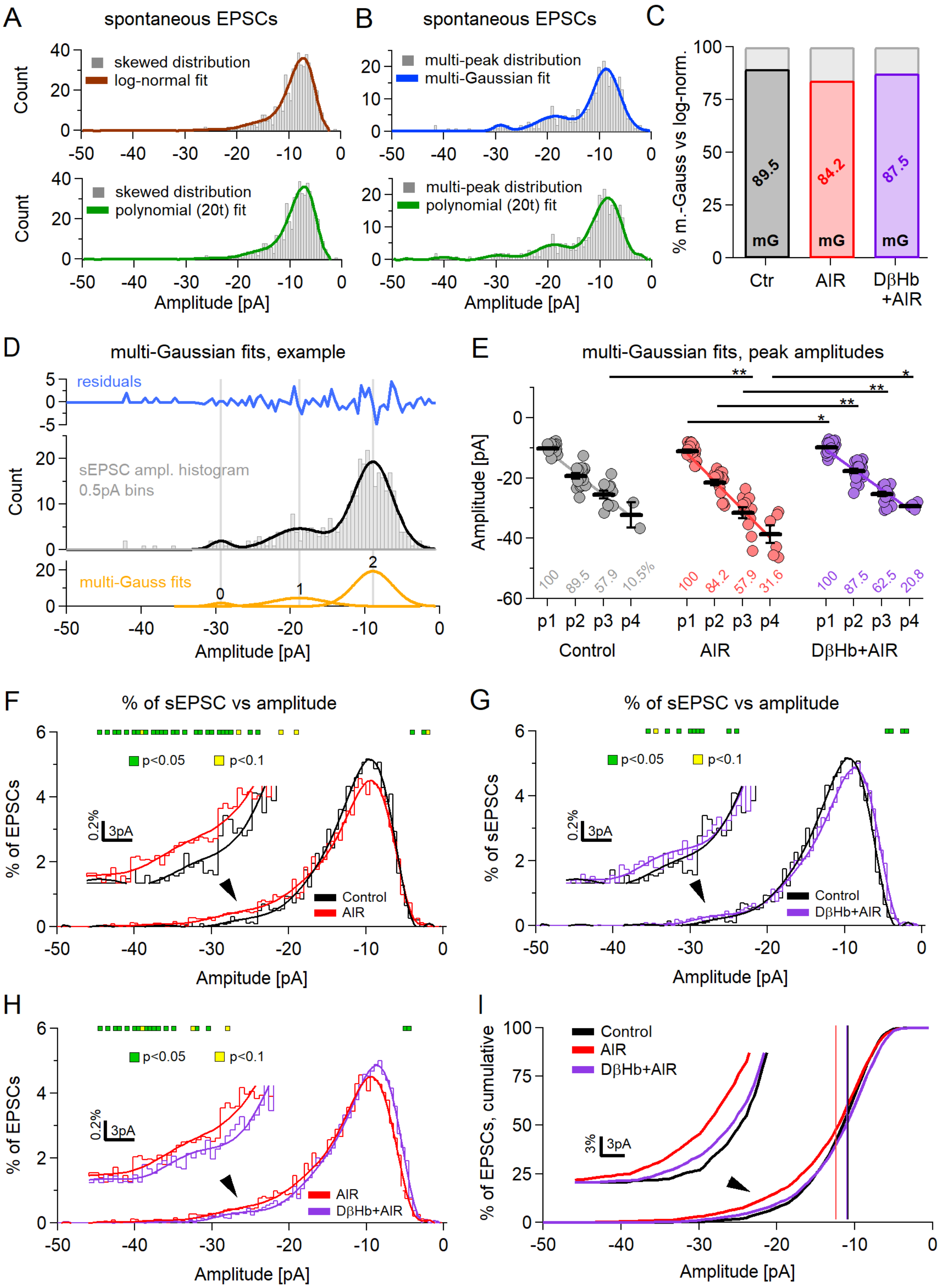
A
spontaneous EPSCs
skewed distribution
log-normal fit
polynomial (20t) fit
Count
Amplitude [pA]
B
spontaneous EPSCs
multi-peak distribution
multi-Gaussian fit
polynomial (20t) fit
C
% m.-Gauss vs log-norm.
89.5
84.2
87.5
mG
Ctr
AIR
DβHb +AIR
D
multi-Gaussian fits, example
residuals
sEPSC ampl. histogram 0.5pA bins
multi-Gauss fits
E
multi-Gaussian fits, peak amplitudes
Amplitude [pA]
p1 p2 p3 p4
Control
AIR
DβHb+AIR
F
% of sEPSC vs amplitude
p<0.05
p<0.1
% of EPSCs
Control
AIR
G
% of sEPSC vs amplitude
% of sEPSCs
DβHb+AIR
H
I
% of EPSCs, cumulative

**Figure S3: Detailed analysis of the sEPSC amplitude distributions reveals an increase in the amplitudes and number of multiquantal sEPSCs.**

**A)** An example of a skewed distribution of sEPSC amplitudes. Skewed distributions have a single peak and are best fitted by a log-normal function (top, brown) or multiterm polynomial (bottom, green). **B)** An example of a multipeaked distribution of sEPSC amplitudes. Multipeak distributions can be fitted by a series of Gaussian functions (navy blue). Multi-Gaussian functions tend to fit better than multiterm polynomials (bottom, green). The peaks of the distribution are expected to correlate to the quanta of neurotransmitter release. **C)** Ratio of multipeak distributions (colored) to skewed distributions (gray) in the experimental groups. Control in black, AIR in red, D-βHb + AIR in purple. **D)** An example of the multi-Gaussian fit with postfit residuals (blue), the fitted distribution (gray), fitted function (black), and 3 individual Gaussians comprising the fitting function (orange). The gray vertical lines denote the peaks of the Gaussian fits. **E)** Scatter plots of the sEPSC multipeak amplitudes compared among the experimental groups. Each colored circle represents a value recorded in a single cell at a specific distribution peak (p1 = 1st peak; p2 = 2nd peak; etc.). Control in gray, AIR in light red, 1 mM D-βHb + AIR in purple. Vertical black bars represent the mean ± SEM. *$p<0.05$, Control n=19, N=19; AIR n=19, N=14; 1 mM D-βHb + AIR, n=24, N=16. The values below the means denote the percentage of cells in which a specific distribution peak could be detected. **F)** Comparison of the averaged and normalized sEPSC amplitude distributions in the Control and AIR groups, binned every 0.5 pA. Note that averaging smooths out multiple peaks of the distributions. The inset shows a magnified part of both distributions around the point with the largest difference. The green squares mark significant differences ($p<0.05$), and the yellow squares mark differences close to significance ($p<0.1$). **G)** As in F), for the Control and D-βHb + AIR distributions. **H)** As in F), for the AIR and D-βHb + AIR distributions. **I)** Distributions from F-H presented as cumulative distributions. The inset is a magnified part of the distributions around the point with the largest difference. The vertical lines mark the average sEPSC amplitudes, as in Fig. 7E.

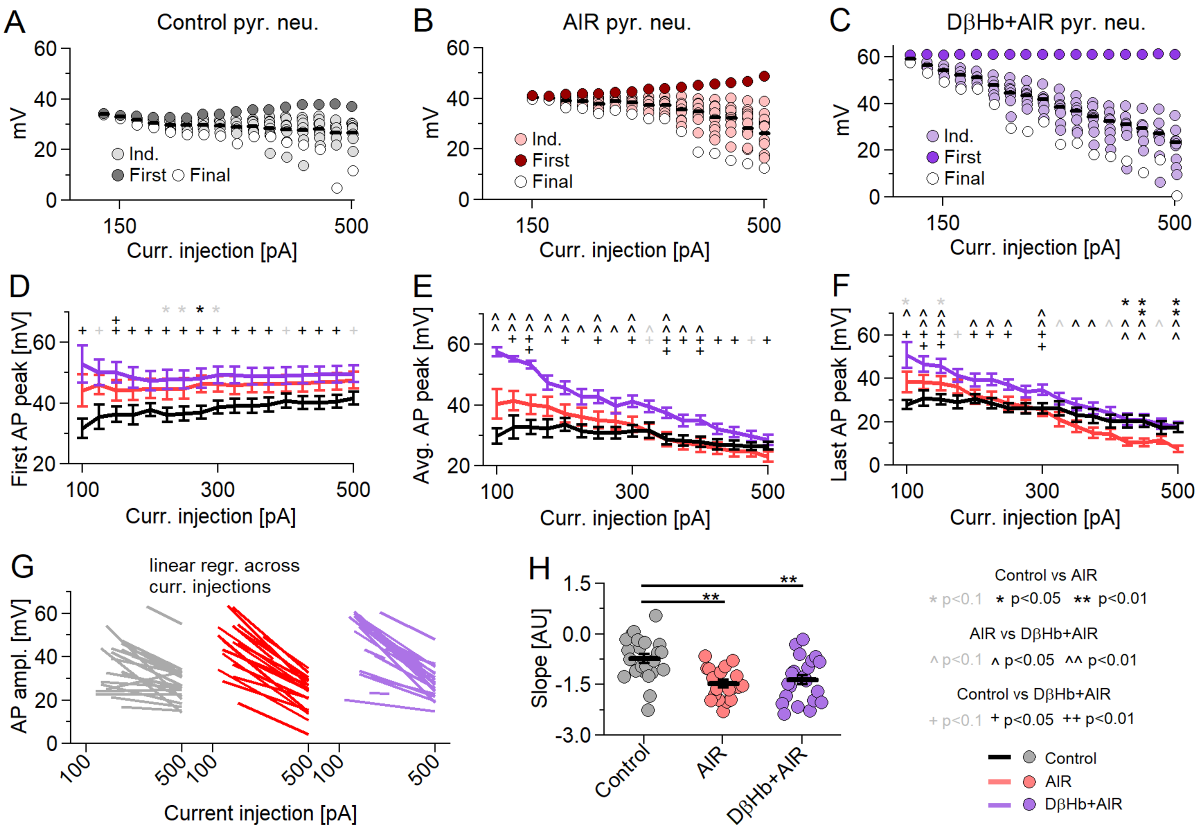
A
Control pyr. neu.
B
AIR pyr. neu.
C
DβHb+AIR pyr. neu.
Ind.
First
Final
mV
Curr. injection [pA]
D
First AP peak [mV]
E
Avg. AP peak [mV]
F
Last AP peak [mV]
G
linear regr. across
curr. injections
AP ampl. [mV]
Current injection [pA]
H
Slope [AU]
Control
AIR
DβHb+AIR
Control vs AIR
* p<0.1 * p<0.05 ** p<0.01
AIR vs DβHb+AIR
^ p<0.1 ^ p<0.05 ^^ p<0.01
Control vs DβHb+AIR
+ p<0.1 + p<0.05 ++ p<0.01
Control
AIR
DβHb+AIR

**Figure S4: Under AIR and D-βHb + AIR conditions, APs show steeper decreases in amplitudes during current injections triggering multiple AP responses.**

**A)** Representative example of AP overshoot amplitudes recorded in response to 21 +25 pA square current injections into a representative control pyramidal neuron. Individual AP amplitudes are in gray, the first AP per injection is in dark gray, the last AP per injection is in white, and the means of all APs per injection are represented by the black bar. **B)** The same as A) but for a representative pyramidal neuron recorded under AIR conditions. **C)** The same as A) but for a representative pyramidal neuron recorded under DβB + AIR conditions. **D)** Group means ± SEMs of the first AP overshoot amplitudes during 21 Δ+25 pA square current injections. * Control vs. AIR, + Control vs. D-βHb + AIR, ^ AIR vs. D-βHb + AIR. *,+,^ in gray $p<0.1$; *,+,^ in black $p<0.05$; **,++,^^ in black $p<0.01$; ANOVA. Control n=23, N=20; AIR n=20, N=14; 1 mM D-βHb + AIR n=22, N=16. **E)** Group means ± SEMs of the average AP overshoot amplitudes during 21 Δ+25 pA square current injections (as in Fig. 8I). N and n are identical to D). **F)** Group means ± SEMs of the final AP overshoot amplitudes during 21 Δ+25 pA square current injections. N and n are identical to D). **G)** Linear regression fits to the average AP overshoot amplitudes in individual pyramidal neurons in all experimental groups. N and n are identical to D). **H)** The slopes of the linear regression fits, compared among the groups. Each colored circle represents a slope for a single cell. Control in gray, AIR in light red, D-βHb + AIR in purple. Vertical black bars represent the mean ± SEM. **$p<0.01$. N and n are identical to D).

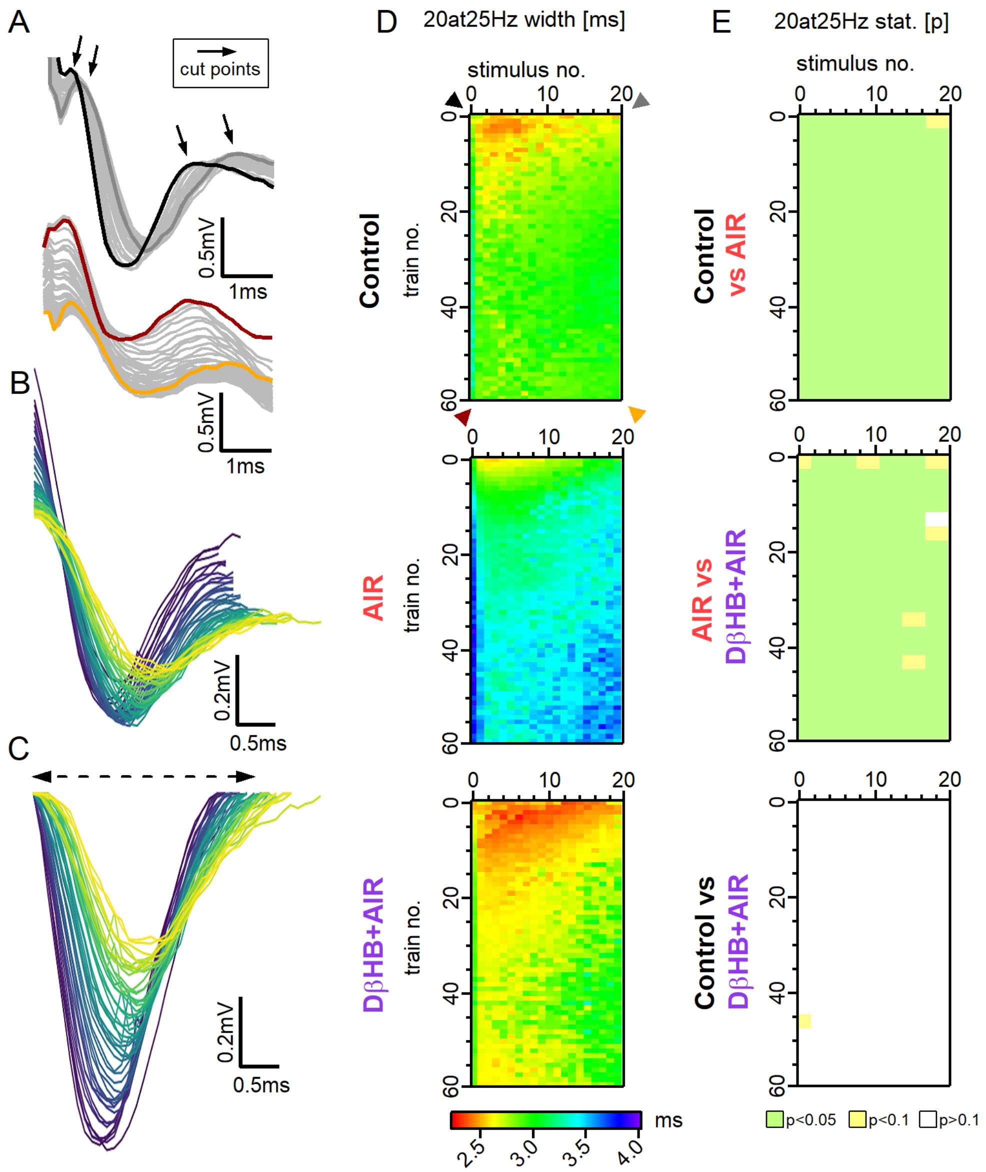
A
cut points
0.5mV
1ms
B
0.2mV
0.5ms
C
D
20at25Hz width [ms]
stimulus no.
train no.
Control
AIR
DβHB+AIR
ms
2.5
3.0
3.5
4.0
E
20at25Hz stat. [p]
Control vs AIR
AIR vs DβHB+AIR
Control vs DβHB+AIR
p<0.05
p<0.1
p>0.1

**Figure S5: AIR causes an increase in the FV widths, while D-βHb, when applied under AIR conditions, recovers and decreases FV widths.**

**A)** Top: Representative examples of the FVs recorded in the control group, at the first stimulus, during stimulation with 20 pulses applied at 25 Hz every 20 s over 20 min. Responses to the first train are shown in black, responses to the last train are shown in dark gray, and other responses are shown in light gray. Bottom: FVs recorded during the same stimulation at the last stimulus (20). Responses to the first train are shown in brown, responses to the last train are shown in orange, and other responses are shown in light gray. The arrows mark the points at which the FVs are cut for subsequent width measurements. **B)** Representative example of 60 FV waveforms extracted from a control train at the first stimulus. **C)** The same FVs, corrected for width analysis. **D)** Heatmaps of mean FV widths recorded over 20 min (60 trains) in the Control, AIR, and 1 mM D-βHb+AIR groups and the significance of the differences. The areas are color-coded as a visible light spectrum (rainbow LUT), with a range of 2.2 (red) to 4.0 ms (violet). Control n=24, N=24; AIR n=22, N=19; D-βHb + AIR n=12, N=10. **E)** P value maps of significant differences among the Control, AIR and 1 mM D-βHb+AIR groups. $p>0.1$ in white; $p<0.1$, $>0.05$ in yellow; $p<0.05$ in green. Note that the comparison was performed based on 3x3 averages.

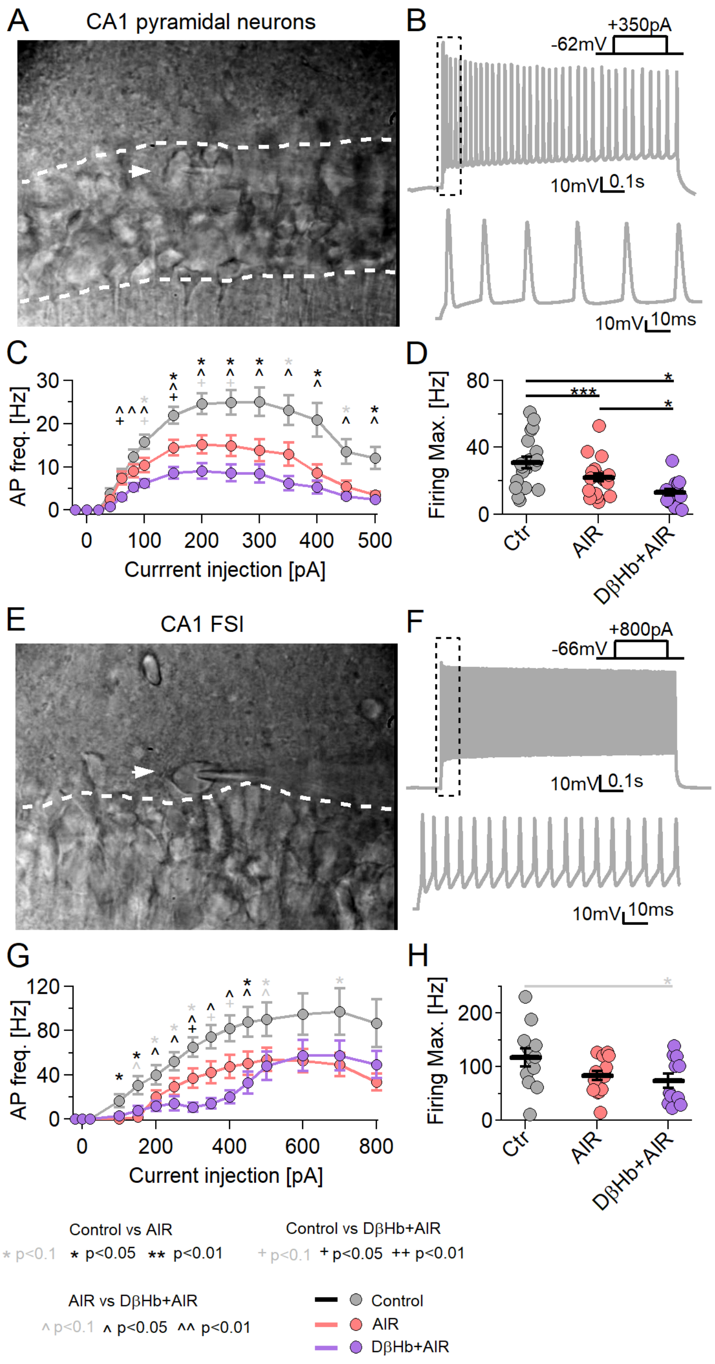
A
CA1 pyramidal neurons
B
+350pA
-62mV
10mV
0.1s
10mV
10ms
C
AP freq. [Hz]
Currrent injection [pA]
D
Firing Max. [Hz]
Ctr
AIR
DβHb+AIR
E
CA1 FSI
F
+800pA
-66mV
10mV
0.1s
10mV
10ms
G
AP freq. [Hz]
Current injection [pA]
H
Firing Max. [Hz]
Ctr
AIR
DβHb+AIR
Control vs AIR
* p<0.1 * p<0.05 ** p<0.01
Control vs DβHb+AIR
+ p<0.1 + p<0.05 ++ p<0.01
AIR vs DβHb+AIR
^ p<0.1 ^ p<0.05 ^^ p<0.01
Control
AIR
DβHb+AIR

**Figure S6: In the absence of inhibition and synaptic inputs, neuronal firing is negatively affected by AIR and further decreases under D-βHb administration.**

**A)** Representative DIC image of a hippocampal CA1 area acquired under 60x magnification. The white dashed lines highlight the pyramidal layer. The white arrowhead indicates a patch-clamped pyramidal neuron. **B)** Top: Representative firing pattern of a CA1 control pyramidal neuron injected with 350 pA current. $V_{rest.}$ = -62 mV. Bottom: The first 100 µs of the injection. **C)** Input–output curves for the CA1 pyramidal neurons in the experimental groups. Each circle represents the group mean ± SEM. * Control vs. AIR, + Control vs. 1 mM D-βHb + AIR, ^ AIR vs. 1 mM D-βHb + AIR. *,+,^ in black p<0.05; **,++,^^ in black p<0.01; *,+,^ in gray p<0.1; ANOVA, Control n=20, N=4; AIR n=20, N=3; 1 mM D-βHb + AIR, n=14, N=2. **D)** Maximum firing rate of the pyramidal neurons. The black bars represent group means ± SEMs. Circles represent the firing rates of individual neurons. * p<0.05, ** p<0.01, p ***<0.001. **E)** Representative DIC image of a hippocampal CA1, as in A). The white arrowhead indicates a patch-clamped FSI interneuron outside the pyramidal layer. **F)** Representative firing pattern of a CA1 control FSI interneuron injected with 800 pA current. Vrest. = -66 mV. Bottom: The first 100 µs of the injection. **G)** Input–output curves for the CA1 FSI in the experimental groups. Each circle represents the group mean ± SEM. Statistical tests and labels are identical to those in C). Control n=12, N=4; AIR n=14, N=4; 1 mM D-βHb + AIR, n=11, N=2. **H)** Maximum firing rate of the FSI in G). Statistical tests and labels are identical to those in C). No significant differences were observed (ANOVA, p=0.068).

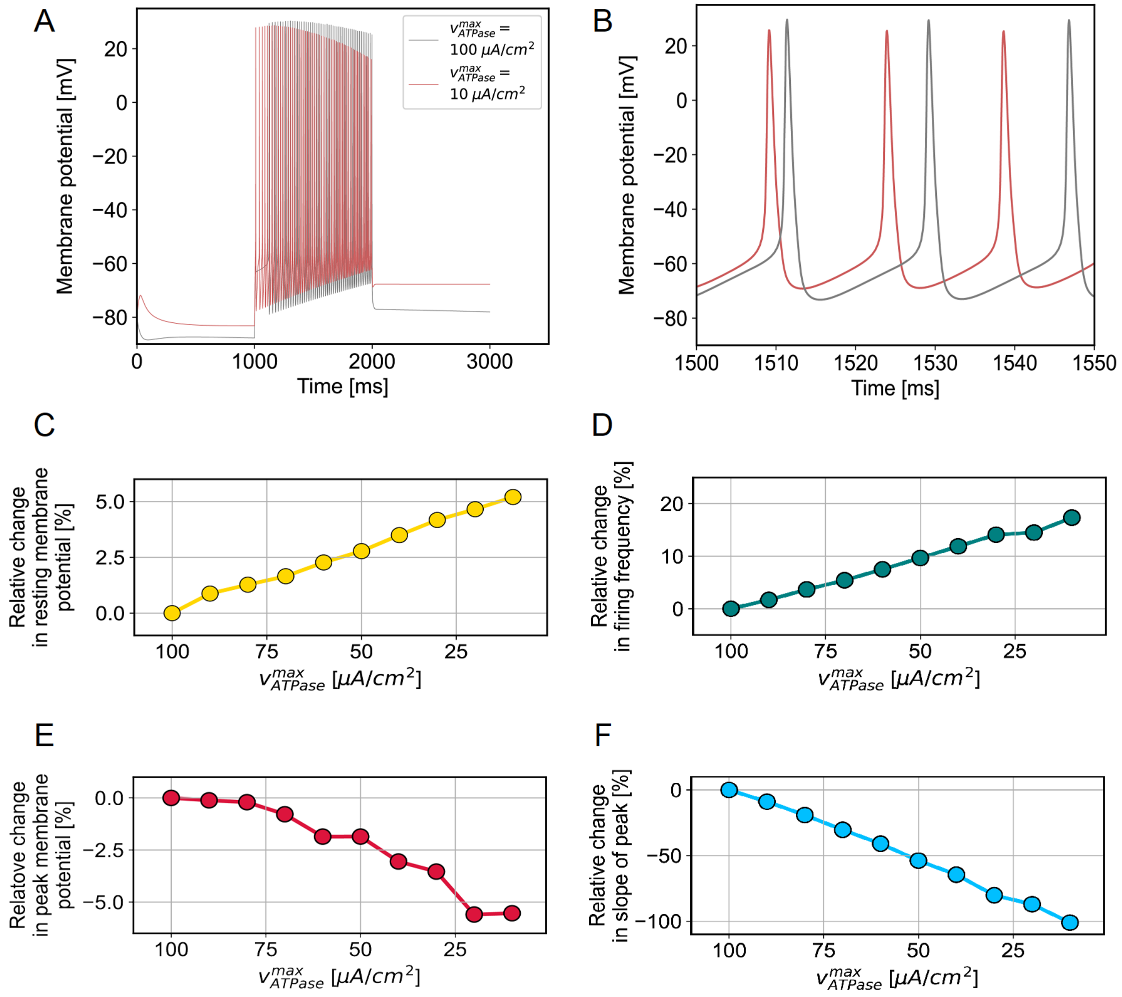


**Figure S7: Impairments in $Na^{+}/K^{+}$ ATPase activity are predicted to have significant effects on neuronal dynamics.**

**A)** Examples of simulated spike trains in response to a 1 s stimulus are shown overlaid at the two extremes of the investigated $v_{ATPase}^{max}$ regime. **B)** Parts of the same spike trains shown with finer temporal resolution. **C-F)** Trends in the resting membrane potential (C), firing frequency (D), peak potential of the first spike (E) and slope of the decline in the peak membrane potential over subsequent spikes (F) are shown as a function of $v_{ATPase}^{max}$, with smaller values on the x-axis representing declining ATPase activity.

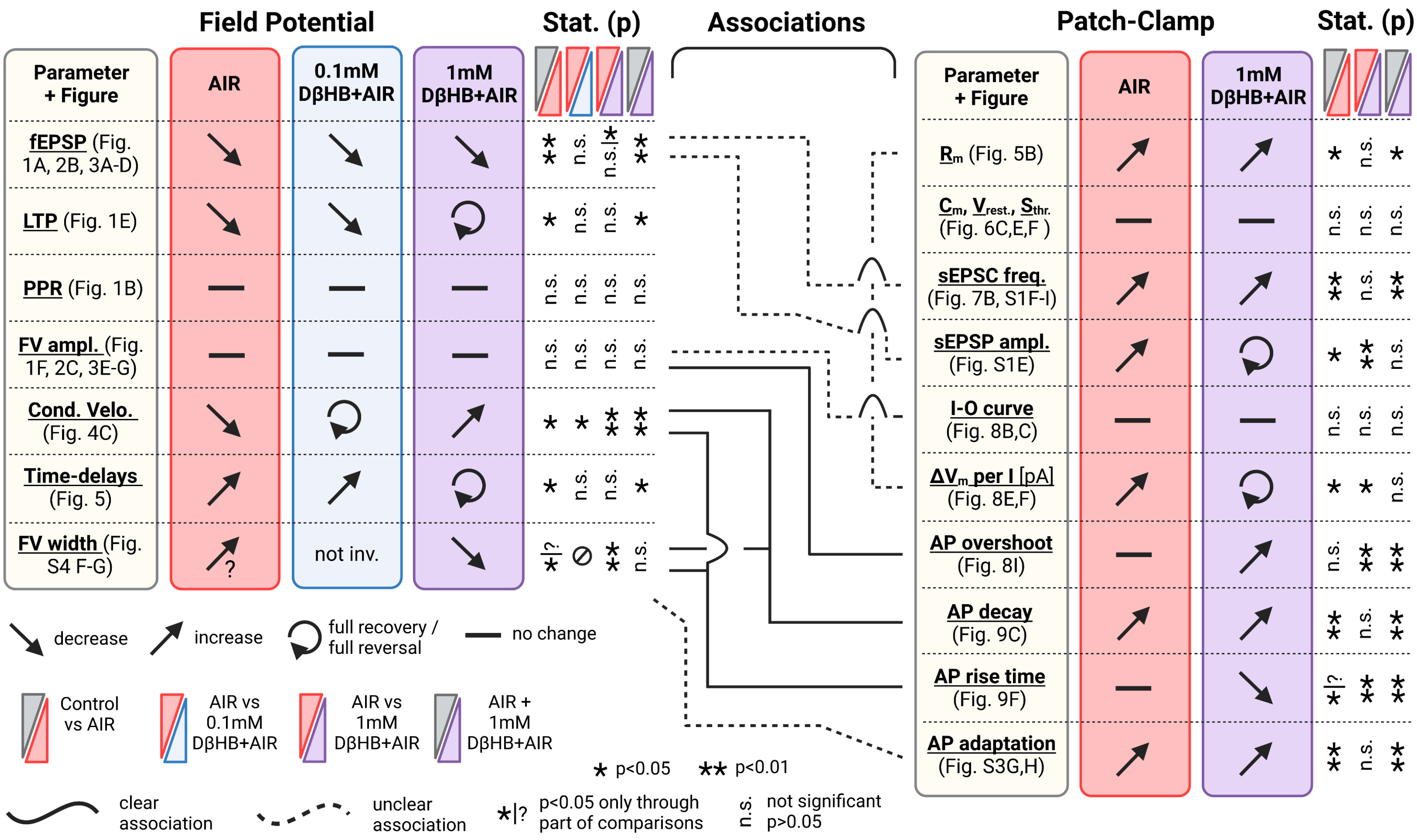

Field Potential
Stat. (p)
Associations
Patch-Clamp
Stat. (p)
Parameter + Figure
AIR
0.1mM DβHB+AIR
1mM DβHB+AIR
fEPSP (Fig. 1A, 2B, 3A-D)
LTP (Fig. 1E)
PPR (Fig. 1B)
FV ampl. (Fig. 1F, 2C, 3E-G)
Cond. Velo. (Fig. 4C)
Time-delays (Fig. 5)
FV width (Fig. S4 F-G)
not inv.
Rm (Fig. 5B)
Cm, Vrest., Sthr. (Fig. 6C,E,F )
sEPSC freq. (Fig. 7B, S1F-I)
sEPSP ampl. (Fig. S1E)
I-O curve (Fig. 8B,C)
ΔVm per I [pA] (Fig. 8E,F)
AP overshoot (Fig. 8I)
AP decay (Fig. 9C)
AP rise time (Fig. 9F)
AP adaptation (Fig. S3G,H)
n.s.
decrease
increase
full recovery / full reversal
no change
Control vs AIR
AIR vs 0.1mM DβHB+AIR
AIR vs 1mM DβHB+AIR
AIR + 1mM DβHB+AIR
* p<0.05
** p<0.01
clear association
unclear association
*|? p<0.05 only through part of comparisons
n.s. not significant p>0.05

**Figure S8: Overall summary of key findings.**

Left: Most important results of the field potential recordings. Right: Most important results of the patch-clamp experiments. The black solid lines link field potential results with their counterparts during patch-clamp experiments. The dashed lines link field potential results with their most likely patch-clamp counterparts, although the associations are less clear. Figure prepared with BioRender.

Supplementary Methods

1.1 *Multi-Gaussian fitting [Figure S3]*

To create histograms of the sEPSC amplitudes, we measured the amplitudes at the peak for each individual waveform using custom-written macros (see Methods, 5.6). Then, we created histograms of the amplitudes with Igor Pro's build-in histogram function by binning the amplitudes into 100 0.5 pA bins ranging from -50 to 0 pA. Next, we imported the histograms into the Multipeak Fitting 2.16 built-in package in Igor Pro. We started the analysis by using the "Auto-locate Peaks Now" option, which utilizes an automatic peak-finding algorithm to search for peaks by finding maxima in the smoothed second derivative of the data. The algorithm automatically estimates the noise level in the data and an optimal smoothing factor. If the initial guess appeared correct, the data were fitted with the multi-Gaussian function suggested by the software. If the fits appeared erroneous, the smoothing factor, widths, and locations of the Gaussian peaks were corrected manually until the best possible fit was achieved. If no corrections provided a good fit, the distributions were fitted by a single Gaussian to capture the largest peak.

1.2 *Whole-cell patch-clamp recordings under abolished AMPA/kainatergic and GABAergic inputs. [Figure S5]*

Acute coronal brain slices (300 μm thick) containing the hippocampus were prepared from male C57BL/6J mice (6 to 8 weeks old; Charles River). The brain was rapidly extracted and soaked in ice-cold (ice slush) oxygenated buffer (in mM): 98 N-methyl-*D*-glucamine (NMDG), 25 D-glucose, 30 $NaHCO_3$, 20 HEPES, 5 Na-*L*-ascorbate, 2 thiourea, 2 ethyl pyruvate, 2.5 KCl, 1.25 $NaH_2PO_4$, 10 $MgSO_4$, 0.5 $CaCl_2$ and 12 N-acetyl-*L*-cysteine (adjusted to pH = 7.4) for cutting. Acute brain slices were prepared with a VT1200S vibratome (Leica Biosystems), with cutting parameters set to 0.08 mm/s speed and 1.2 mm vibration amplitude. Then, 10 to 12 slices containing the hippocampus were collected and left to recover for 10 minutes in a 32 °C NMDG solution in an interface chamber (Brain Slice Keeper 5, Scientific Systems Design). Slices were individually timed to ensure the precise timing of this step. Slices were then moved to a second interface chamber containing HEPES holding solution at RT containing (in mM): 90 NaCl, 25 *D*-glucose, 30 $NaHCO_3$, 20 HEPES, 3.5 NaOH, 2 $CaCl_2$, 2 $MgSO_4$, 1.25 $NaH_2PO_4$, 2.5 KCl, 2 thiourea, 5 Na-*L*-ascorbate, and 5 ethyl pyruvate. The slices were then left to recover for an additional 50 minutes. The total recovery time was at least 60 min after cutting.

Slices were transferred to a recording chamber continuously perfused with recording aCSF at a rate of 3-5 ml/min at RT. Whole-cell recordings were performed using a software-controlled MultiClamp 700B amplifier and a Digidata 1440A digitizer (Molecular Devices, USA). Recording pipettes were pulled from borosilicate glass capillaries (WPI) with a resistance of 3-6 MΩ and filled with an intracellular solution containing (in mM): 135 K-gluconate, 3 KCl, 3 $MgCl_2$, 10

HEPES, 0.5 EGTA, 4 $Na_2ATP$, 0.3 $Na_3GTP$, 10 Na-phosphocreatine, pH adjusted to 7.3 with 1 M KOH and osmolarity adjusted to ~290 ± 5 mOsm. Slices were visualized with infrared optics using an upright microscope equipped with differential interference contrast (IR-DIC) using a 4x objective. Pyramidal neurons and fast-spiking interneurons (FSIs) in the CA1 region were visualized with a 60x immersion objective and identified by their morphology and electrophysiological properties (i.e., presence of a voltage "sag" with hyperpolarizing current step injections and specific pattern of AP firing with depolarizing current step injections in current-clamp mode).

Slices were incubated in parallel with aCSF in 0.1% DMSO (control), 0.1 mM indinavir, or 0.1 mM indinavir + 1 mM D-βHb for at least 45 minutes before the recordings, which were obtained directly in the patch chamber over a period of no more than 3 hours, with the inclusion of additional synaptic blockers, namely, 10 μM NBQX and 10 μM gabazine, which were also present in the aCSF throughout the recording. A maximum of 4 neurons per slice were recorded. The mean incubation times were 93 min for the control group and 95 min for the indinavir group.

After a short period of stabilization (3 minutes) in the whole-cell configuration, neurons were recorded without any current injection (the holding was not set within a specific range) in current-clamp mode. The protocol for pyramidal neurons consisted of 3-second sweeps with 1-second pulses of from -20 to 100 pA in steps of 20 and then to 500 pA in steps of 50. For fast-spiking interneurons, we used 3-second sweeps with 1-second pulses from -20 to 100 pA in steps of twenty, to 500 pA in steps of 50, and then to 800 pA in steps of 100. Each group of pulses was separated by 15 seconds, and the results of 3 consecutive runs were averaged.

All data acquisition was performed using pCLAMP software (Molecular Devices, USA).

### 1.3 *Hodgkin-Huxley model [Figure S7]*

Our computational simulations investigated the effects of decreased $Na^+/K^+$ ATPase activity utilizing a model primarily based on a previously validated CA1 neuron model [66]. This model builds upon the original Hodgkin-Huxley model but has greater specificity to various voltage-gated ion channels, including separate $Ca^{2+}$ and $Na^+$ channels and various types of $K^+$ channels. We combined this framework with additional components specific to $Na^+/K^+$ ATPase, which were adapted from [67]. These additions involved kinetic equations for concentrations and reversal potentials specific to $Na^+$ and $K^+$ and a dependence on ATPase activity. To investigate the effects of different $Na^+/K^+$ ATPase activity rates, we compared the kinetic parameter $v_{ATPase}^{max}$ between 10 μA/cm$^2$ and 100 μA/cm$^2$ in increments of 10 μA/cm$^2$. $v_{ATPase}^{max}$ determines $Na^+/K^+$ ATPase activity in the following manner:

$$I_{pump} = v_{ATPase}^{max} \left(1 + \frac{K_M^{Na}}{[N]_i}\right)^{-3} \left(1 + \frac{K_M^K}{[K]_e}\right)^{-2},$$

where $[Na]_i$ and $[K]_e$ are ion concentrations that vary over time, and $K_M^{Na}$ and $K_M^K$ are Michaelis–Menten constants. The extension with pump dynamics necessitated specification of a volume and surface area for the CA1 neuron. Our modeled neuron had a volume of 2000 $\mu m^3$ [68], and we assumed that the neuron had a smooth spherical surface and an active surface area of 800 $\mu m^2$. The temperature was set at 24.5 °C to closely match our experimental conditions. During each simulation, the neuron was left at rest for 1 s to reach equilibrium and then stimulated for 1 s at 3.125 $\mu A/cm^2$, equivalent to 250 pA. The model was implemented in the Julia programming language and simulated with the DifferentialEquiations.jl library (https://github.com/SciML/DifferentialEquations.jl).

To evaluate the effects of diminished $Na^+/K^+$ ATPase activity, we extracted multiple features from the simulated spike trains. The resting state potential (mV) was quantified as the membrane potential at 1 s immediately before stimulation. The firing frequency (Hz) was calculated by dividing the number of APs by the time duration between the first and last evoked spike. The peak membrane potential (mV) was determined from the first AP of the spike train. Finally, we quantified the rate of exhaustion during a spike train by fitting a linear model to the progression of peak potentials over the index number of the corresponding AP. The slope of the fitted line was extracted and labeled as the slope of the peak.

All relevant code is available on GitHub, including all equations and parameters. The code can also be used to reproduce all results and is available at: https://github.com/lcneuro/iAIR_models.
