## Supplemental File for "*D*-*ꞵ*-hydroxybutyrate stabilizes hippocampal CA3-CA1 circuit during acute insulin resistance"

Bartosz Kula *et al.*

**This PDF file includes:**

Figs. S1 to S8

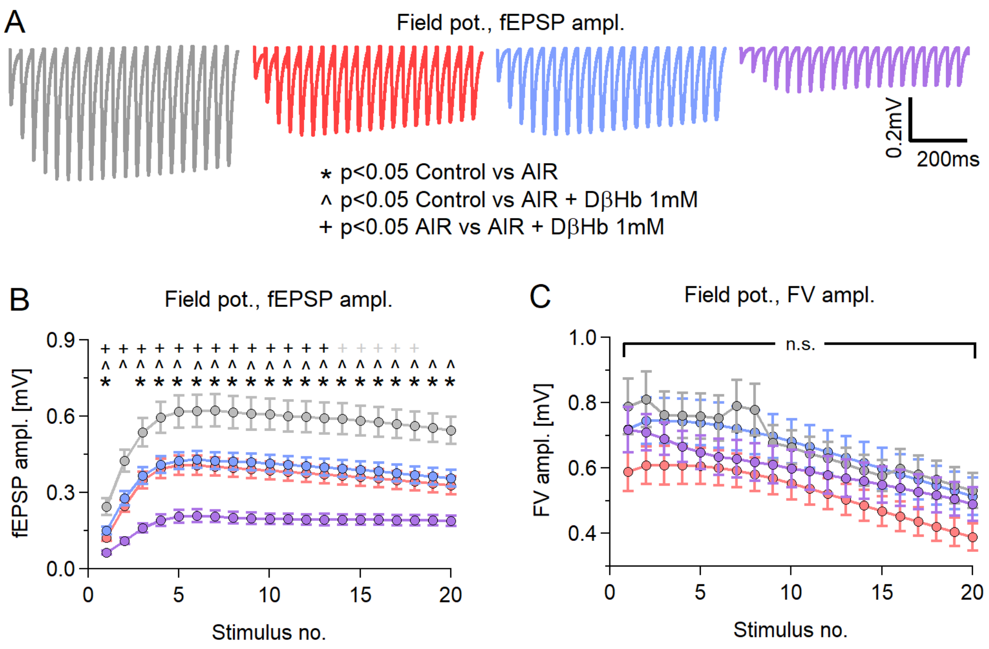


**Figure S1: AIR and D-βHb have the same effects during train and paired stimulation.**

A) Representative examples of averaged fEPSP responses, recorded in each experimental group, to Schaffer collateral stimulation with 20 pulses applied at 25 Hz every 20 s for 20 min, with 280-350 µm distances between electrodes. Control in gray; AIR in red; 0.1 mM D-βHb+AIR in blue; 1 mM D-βHb+AIR in purple. Stimulation artifacts and FVs are removed for clarity. B) Group mean fEPSP amplitudes ± SEM evoked by stimulation, as in A), plotted against the stimulus number. *, +, and ^ indicate statistically significant differences. *<0.05 Control vs. 0.1 mM AIR; #p<0.05 Control vs. 0.1 mM D-βHb+AIR; +p<0.05 Control vs. 1 mM D-βHb+AIR; ^p<0.05 0.1 mM D-βHb+AIR vs. 1 mM D-βHb+AIR; ANOVA. Control, n=24, N=22; AIR, n=16, N=15; 0.1 mM D-βHb+AIR, n=17, N=15; 1 mM D-βHb+AIR, n=14, N=10. C) Group mean FV amplitudes ± SEM recorded during A), plotted against the stimulus number. The labels n and N are identical to (B). There were no significant differences (p=0.091 to 0.34 during the trains, ANOVA).

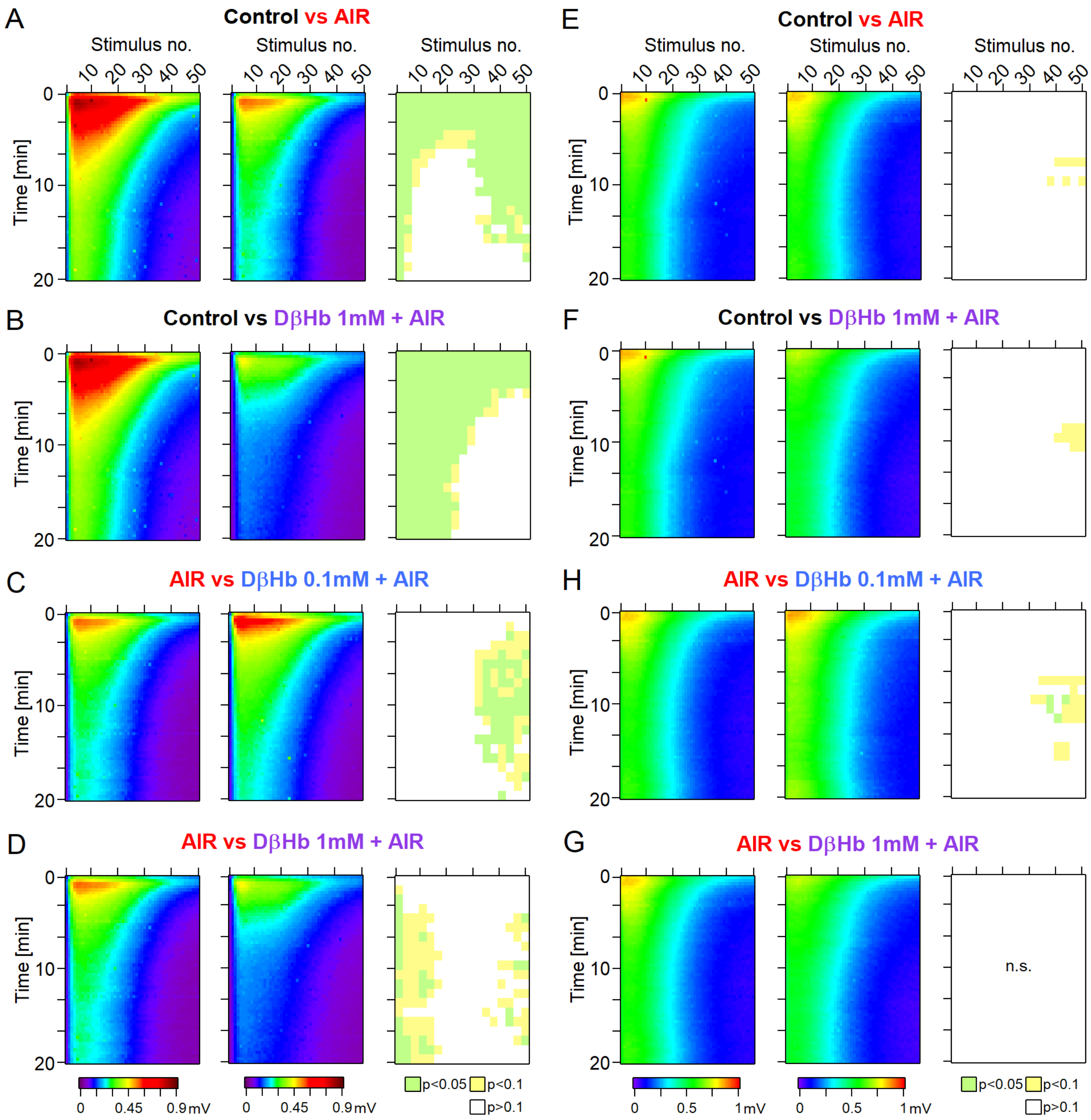


**Figure S2: During overtaxing, nonphysiological stimulation, AIR and D-βHb show comparable effects with physiological stimulation paradigms.**

**A)** Heatmaps of mean fEPSP amplitudes recorded over 60 min in the Control and AIR groups and the significance of the differences. The y-axis represents the time points of the trains of 50 stimuli applied at 25 Hz every 20 s, with 280-350 µm distances between electrodes. The x-axis represents the time points of responses to individual stimuli. fEPSP amplitudes are color-coded as a visible light spectrum in the range of 0.1 mV (brown) and 0 mV (violet). Control, n=24 slices, N=24 mice; AIR, n=21, N=18. The groups were compared as 3 stimuli x 3 time blocks, and the p values are summarized on the right-most graph. **B)** The same as A) for the Control and 1 mM D-βHb+AIR groups. Control n=24, N=24; 1 mM D-βHb+AIR n=12, N=10. **C)** The same as A) for the AIR and 0.1 mM D-βHb+AIR groups. AIR n=21, N=18; 0.1 mM D-βHb+AIR; n=18, N=15. **D)** The same as A) for the AIR and 1 mM D-βHb+AIR groups. AIR n=21, N=18; 1 mM D-βHb+AIR n=12, N=10. **E)** Comparison of the mean FV amplitudes between the Control and AIR groups, as in A). The layout, color-coding, and n and N numbers are identical to A). **F)** The same as E) for the Control and 1 mM D-βHb+AIR groups. The layout, color coding, and n and N numbers are identical to B). **G)** The same as E) for the AIR and 0.1 mM D-βHb+AIR groups. The layout, color coding, and n and N numbers are identical to C). **H)** The same as E) for the AIR and 1 mM D-βHb+AIR groups. The layout, color coding, and n and N numbers are identical to D).

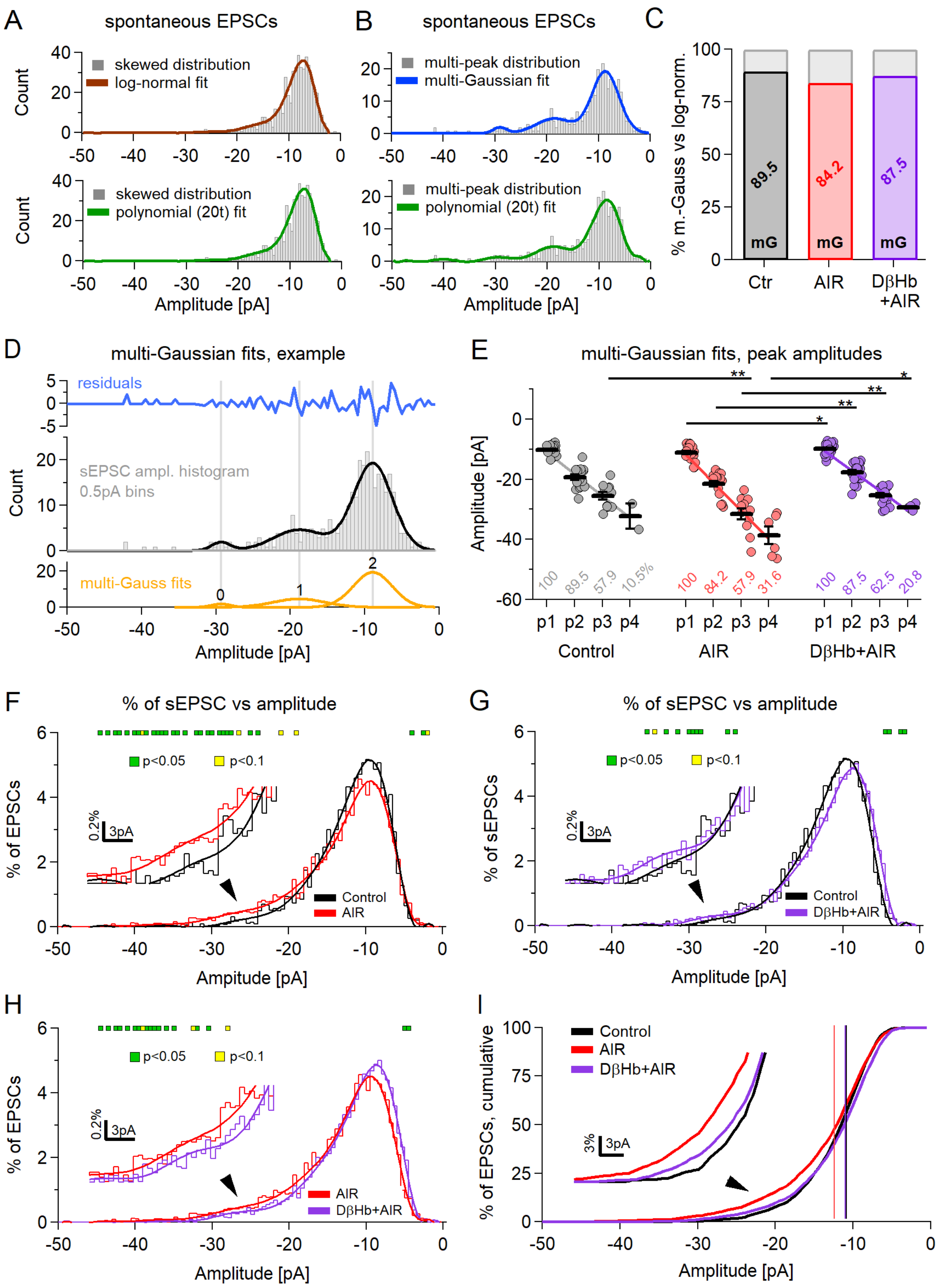

A
spontaneous EPSCs
skewed distribution
log-normal fit
polynomial (20t) fit
Count
Amplitude [pA]
B
spontaneous EPSCs
multi-peak distribution
multi-Gaussian fit
C
% m.-Gauss vs log-norm.
89.5
84.2
87.5
mG
Ctr
AIR
DβHb +AIR
D
multi-Gaussian fits, example
residuals
sEPSC ampl. histogram 0.5pA bins
multi-Gauss fits
E
multi-Gaussian fits, peak amplitudes
Amplitude [pA]
p1 p2 p3 p4
Control
AIR
DβHb+AIR
F
% of sEPSC vs amplitude
p<0.05
p<0.1
% of EPSCs
Control
AIR
Ampitude [pA]
G
% of sEPSC vs amplitude
% of sEPSCs
DβHb+AIR
H
I
% of EPSCs, cumulative

**Figure S3: Detailed analysis of the sEPSC amplitude distributions reveals an increase in the amplitudes and number of multiquantal sEPSCs.**

**A)** An example of a skewed distribution of sEPSC amplitudes. Skewed distributions have a single peak and are best fitted by a log-normal function (top, brown) or multiterm polynomial (bottom, green). **B)** An example of a multipeaked distribution of sEPSC amplitudes. Multipeak distributions can be fitted by a series of Gaussian functions (navy blue). Multi-Gaussian functions tend to fit better than multiterm polynomials (bottom, green). The peaks of the distribution are expected to correlate to the quanta of neurotransmitter release. **C)** Ratio of multipeak distributions (colored) to skewed distributions (gray) in the experimental groups. Control in black, AIR in red, D-βHb + AIR in purple. **D)** An example of the multi-Gaussian fit with postfit residuals (blue), the fitted distribution (gray), fitted function (black), and 3 individual Gaussians comprising the fitting function (orange). The gray vertical lines denote the peaks of the Gaussian fits. **E)** Scatter plots of the sEPSC multipeak amplitudes compared among the experimental groups. Each colored circle represents a value recorded in a single cell at a specific distribution peak (p1 = 1st peak; p2 = 2nd peak; etc.). Control in gray, AIR in light red, 1 mM D-βHb + AIR in purple. Vertical black bars represent the mean ± SEM. *$p<0.05$, Control n=19, N=19; AIR n=19, N=14; 1 mM D-βHb + AIR, n=24, N=16. The values below the means denote the percentage of cells in which a specific distribution peak could be detected. **F)** Comparison of the averaged and normalized sEPSC amplitude distributions in the Control and AIR groups, binned every 0.5 pA. Note that averaging smooths out multiple peaks of the distributions. The inset shows a magnified part of both distributions around the point with the largest difference. The green squares mark significant differences ($p<0.05$), and the yellow squares mark differences close to significance ($p<0.1$). **G)** As in F), for the Control and D-βHb + AIR distributions. **H)** As in F), for the AIR and D-βHb + AIR distributions. **I)** Distributions from F-H presented as cumulative distributions. The inset is a magnified part of the distributions around the point with the largest difference. The vertical lines mark the average sEPSC amplitudes, as in Fig. 7E.

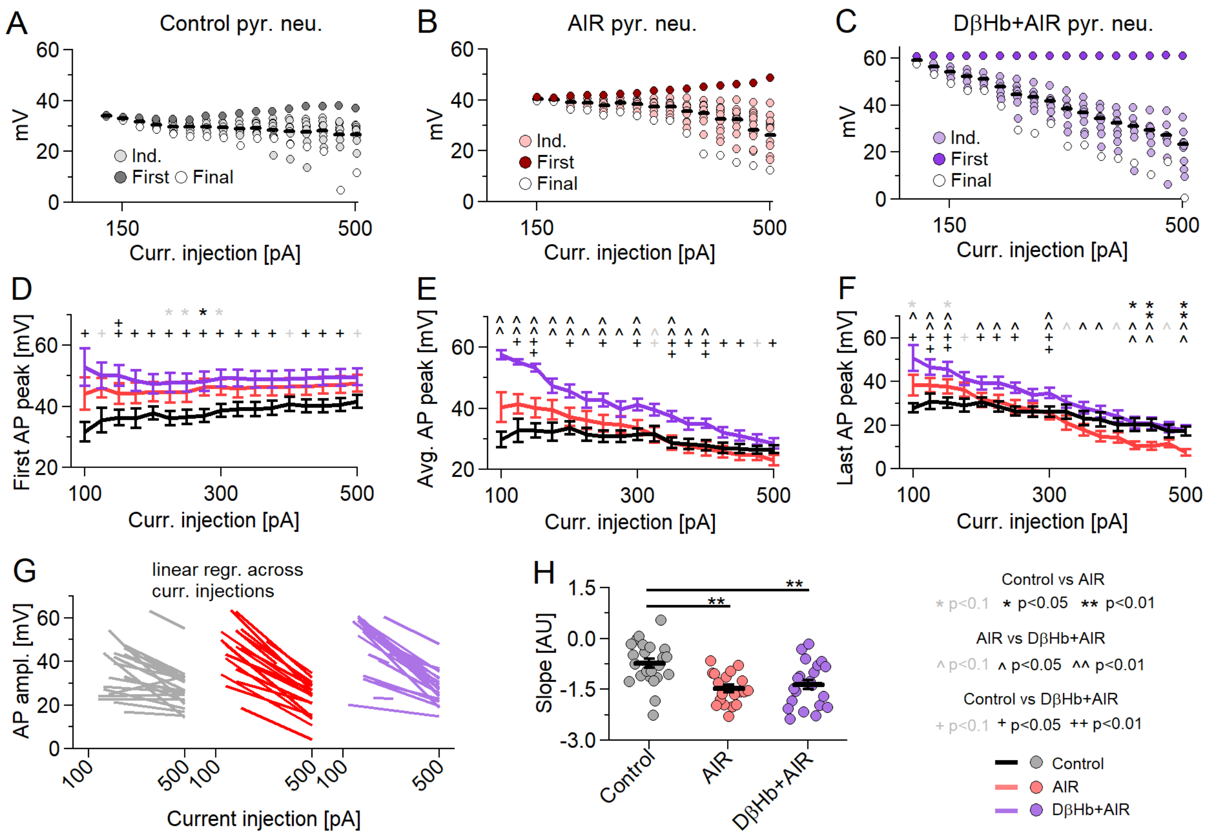


**Figure S4: Under AIR and D-βHb + AIR conditions, APs show steeper decreases in amplitudes during current injections triggering multiple AP responses.**

**A)** Representative example of AP overshoot amplitudes recorded in response to 21 +25 pA square current injections into a representative control pyramidal neuron. Individual AP amplitudes are in gray, the first AP per injection is in dark gray, the last AP per injection is in white, and the means of all APs per injection are represented by the black bar. **B)** The same as A) but for a representative pyramidal neuron recorded under AIR conditions. **C)** The same as A) but for a representative pyramidal neuron recorded under DβB + AIR conditions. **D)** Group means ± SEMs of the first AP overshoot amplitudes during 21 Δ+25 pA square current injections. * Control vs. AIR, + Control vs. D-βHb + AIR, ^ AIR vs. D-βHb + AIR. *,+,^ in gray p<0.1; *,+,^ in black p<0.05; **,++,^^ in black p<0.01; ANOVA. Control n=23, N=20; AIR n=20, N=14; 1 mM D-βHb + AIR n=22, N=16. **E)** Group means ± SEMs of the average AP overshoot amplitudes during 21 Δ+25 pA square current injections (as in Fig. 8I). N and n are identical to D). **F)** Group means ± SEMs of the final AP overshoot amplitudes during 21 Δ+25 pA square current injections. N and n are identical to D). **G)** Linear regression fits to the average AP overshoot amplitudes in individual pyramidal neurons in all experimental groups. N and n are identical to D). **H)** The slopes of the linear regression fits, compared among the groups. Each colored circle represents a slope for a single cell. Control in gray, AIR in light red, D-βHb + AIR in purple. Vertical black bars represent the mean ± SEM. **$p<0.01$. N and n are identical to D).

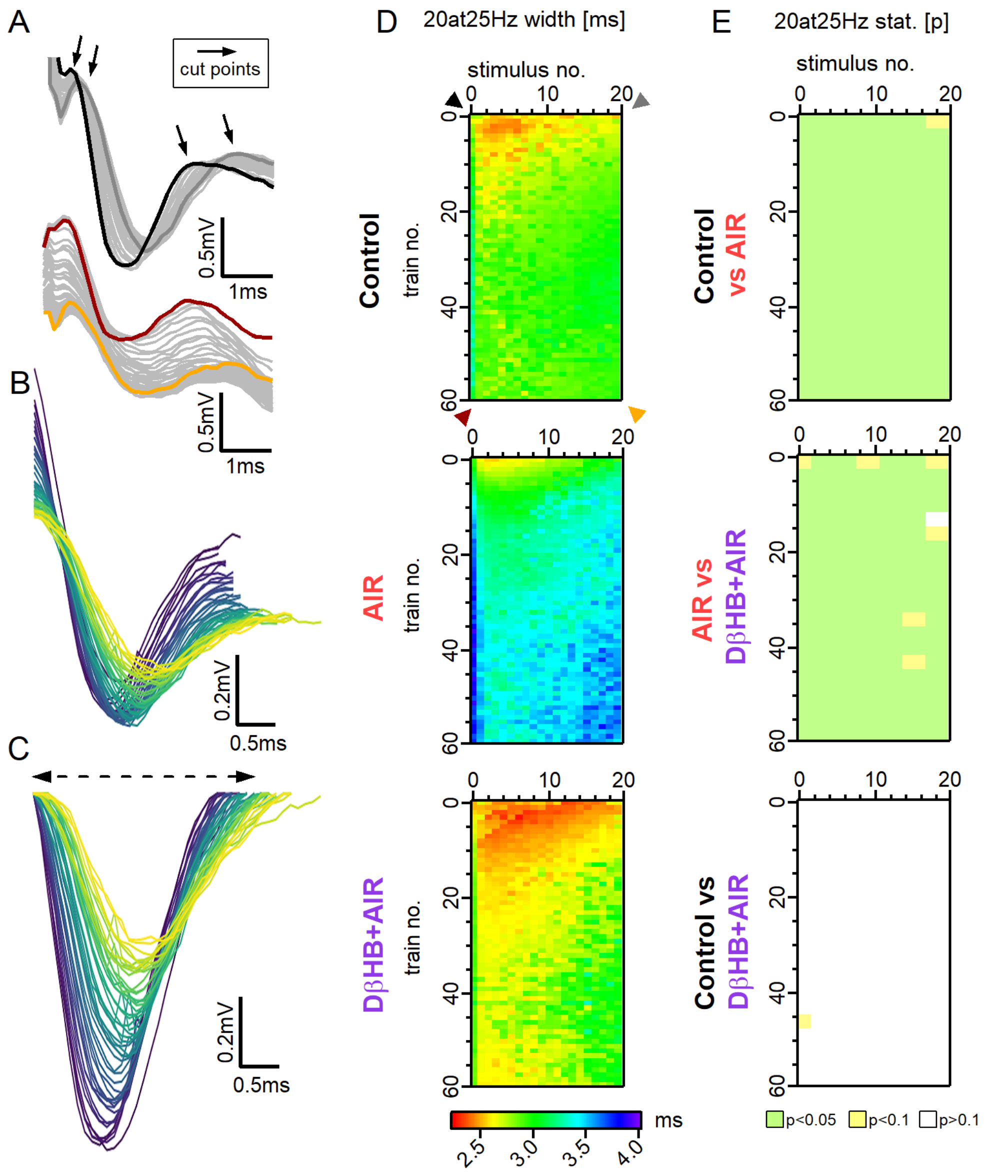


**Figure S5: AIR causes an increase in the FV widths, while D-βHb, when applied under AIR conditions, recovers and decreases FV widths.**

**A)** Top: Representative examples of the FVs recorded in the control group, at the first stimulus, during stimulation with 20 pulses applied at 25 Hz every 20 s over 20 min. Responses to the first train are shown in black, responses to the last train are shown in dark gray, and other responses are shown in light gray. Bottom: FVs recorded during the same stimulation at the last stimulus (20). Responses to the first train are shown in brown, responses to the last train are shown in orange, and other responses are shown in light gray. The arrows mark the points at which the FVs are cut for subsequent width measurements. **B)** Representative example of 60 FV waveforms extracted from a control train at the first stimulus. **C)** The same FVs, corrected for width analysis. **D)** Heatmaps of mean FV widths recorded over 20 min (60 trains) in the Control,

AIR, and 1 mM D-βHb+AIR groups and the significance of the differences. The areas are color-coded as a visible light spectrum (rainbow LUT), with a range of 2.2 (red) to 4.0 ms (violet). Control n=24, N=24; AIR n=22, N=19; D-βHb + AIR n=12, N=10. **E)** P value maps of significant differences among the Control, AIR and 1 mM D-βHb+AIR groups. $p>0.1$ in white; $p<0.1$, $>0.05$ in yellow; $p<0.05$ in green. Note that the comparison was performed based on 3x3 averages.

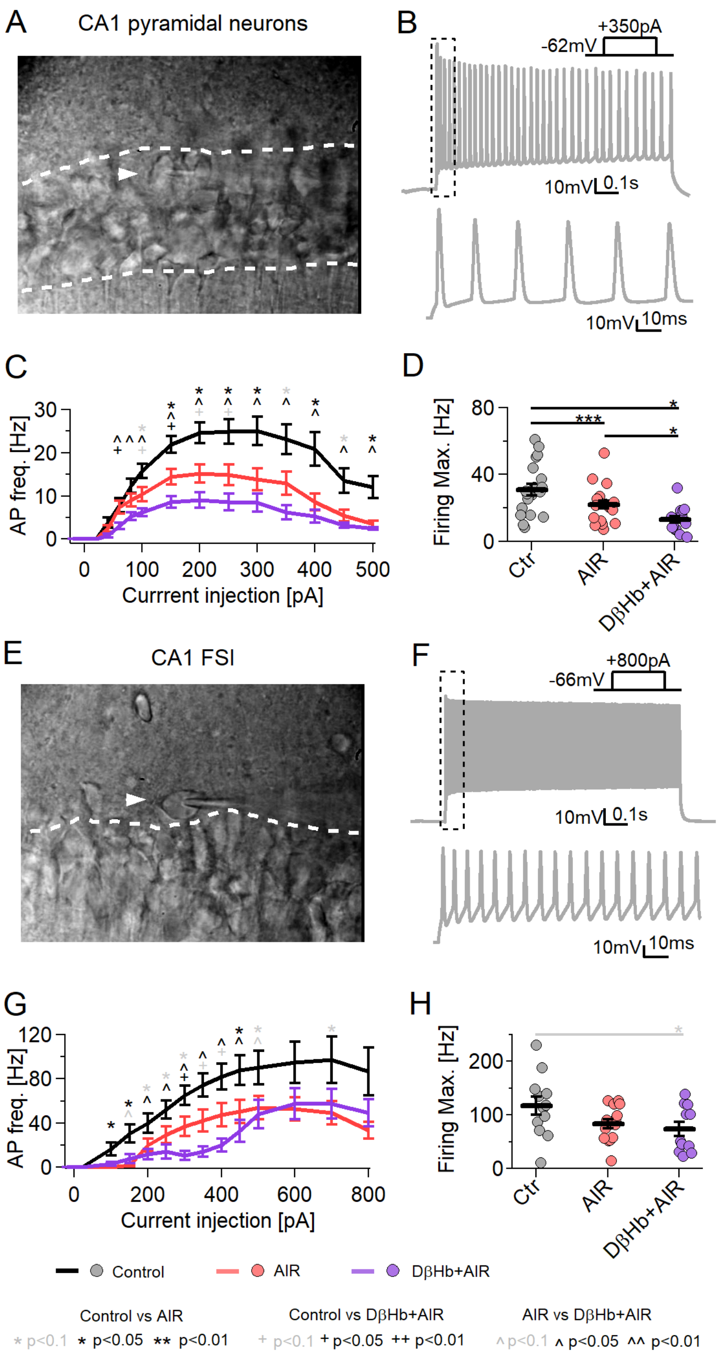


**Figure S6: In the absence of inhibition and synaptic inputs, neuronal firing is negatively affected by AIR and further decreases under D-βHb administration.**

**A)** Representative DIC image of a hippocampal CA1 area acquired under 60x magnification. The white dashed lines highlight the pyramidal layer. The white arrowhead indicates a patch-clamped pyramidal neuron. **B)** Top: Representative firing pattern of a CA1 control pyramidal neuron injected with 350 pA current. $V_{rest.}$ = -62 mV. Bottom: The first 100 µs of the injection. **C)** Input–

output curves for the CA1 pyramidal neurons in the experimental groups. Each circle represents the group mean ± SEM. * Control vs. AIR, + Control vs. 1 mM D-βHb + AIR, ^ AIR vs. 1 mM D-βHb + AIR. *,+,^ in black $p<0.05$; **,++,^^ in black $p<0.01$; *,+,^ in gray $p<0.1$; ANOVA, Control n=20, N=4; AIR n=20, N=3; 1 mM D-βHb + AIR, n=14, N=2. **D)** Maximum firing rate of the pyramidal neurons. The black bars represent group means ± SEMs. Circles represent the firing rates of individual neurons. * $p<0.05$, ** $p<0.01$, p ***<0.001. **E)** Representative DIC image of a hippocampal CA1, as in A). The white arrowhead indicates a patch-clamped FSI interneuron outside the pyramidal layer. **F)** Representative firing pattern of a CA1 control FSI interneuron injected with 800 pA current. Vrest. = -66 mV. Bottom: The first 100 µs of the injection. **G)** Input–output curves for the CA1 FSI in the experimental groups. Each circle represents the group mean ± SEM. Statistical tests and labels are identical to those in C). Control n=12, N=4; AIR n=14, N=4; 1 mM D-βHb + AIR, n=11, N=2. **H)** Maximum firing rate of the FSI in G). Statistical tests and labels are identical to those in C). No significant differences were observed (ANOVA, p=0.068).

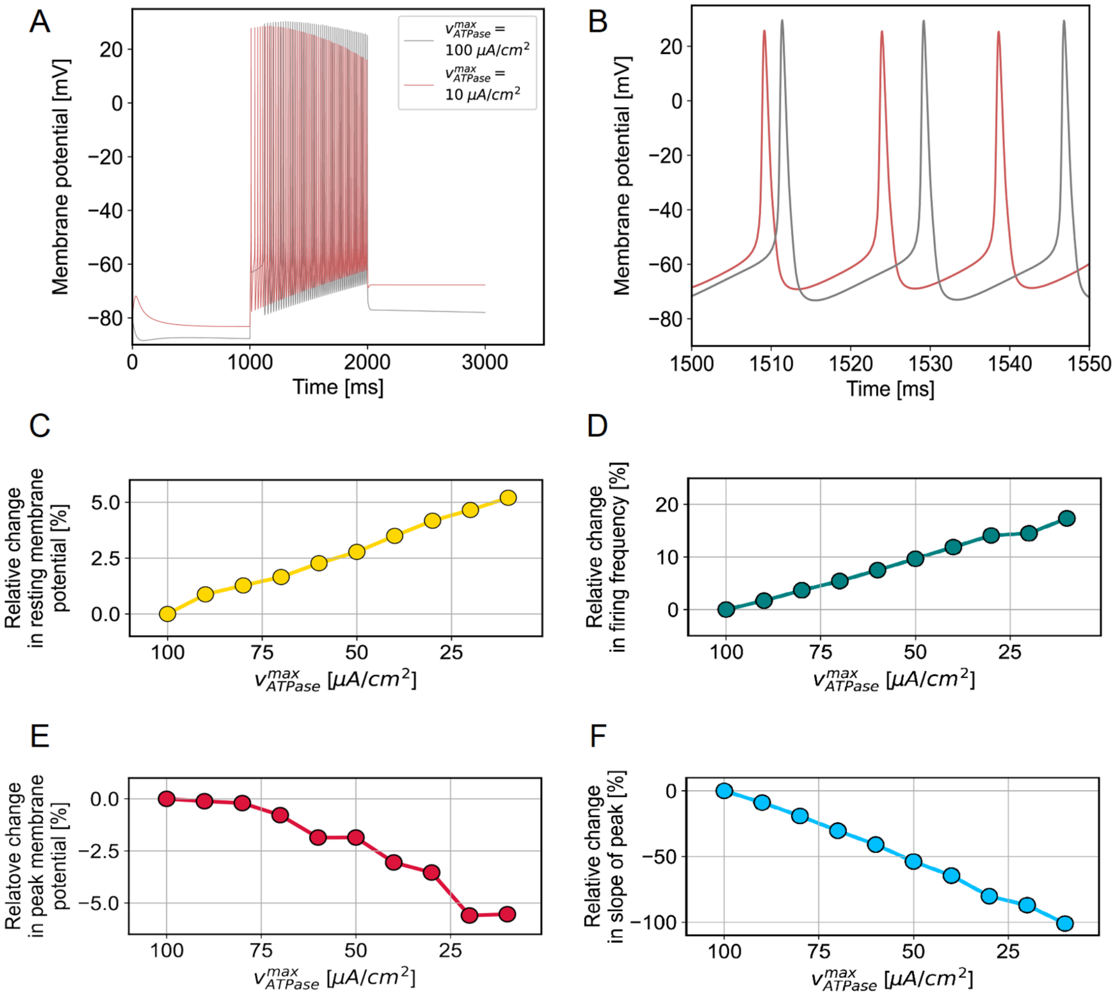


**Figure S7: Impairments in Na$^+$/K$^+$ ATPase activity are predicted to have significant effects on neuronal dynamics.**

**A)** Examples of simulated spike trains in response to a 1 s stimulus are shown overlaid at the two extremes of the investigated $v_{ATPase}^{max}$ regime. **B)** Parts of the same spike trains shown with finer temporal resolution. **C-F)** Trends in the resting membrane potential (C), firing frequency (D), peak potential of the first spike (E) and slope of the decline in the peak membrane potential over subsequent spikes (F) are shown as a function of $v_{ATPase}^{max}$, with smaller values on the x-axis representing declining ATPase activity.

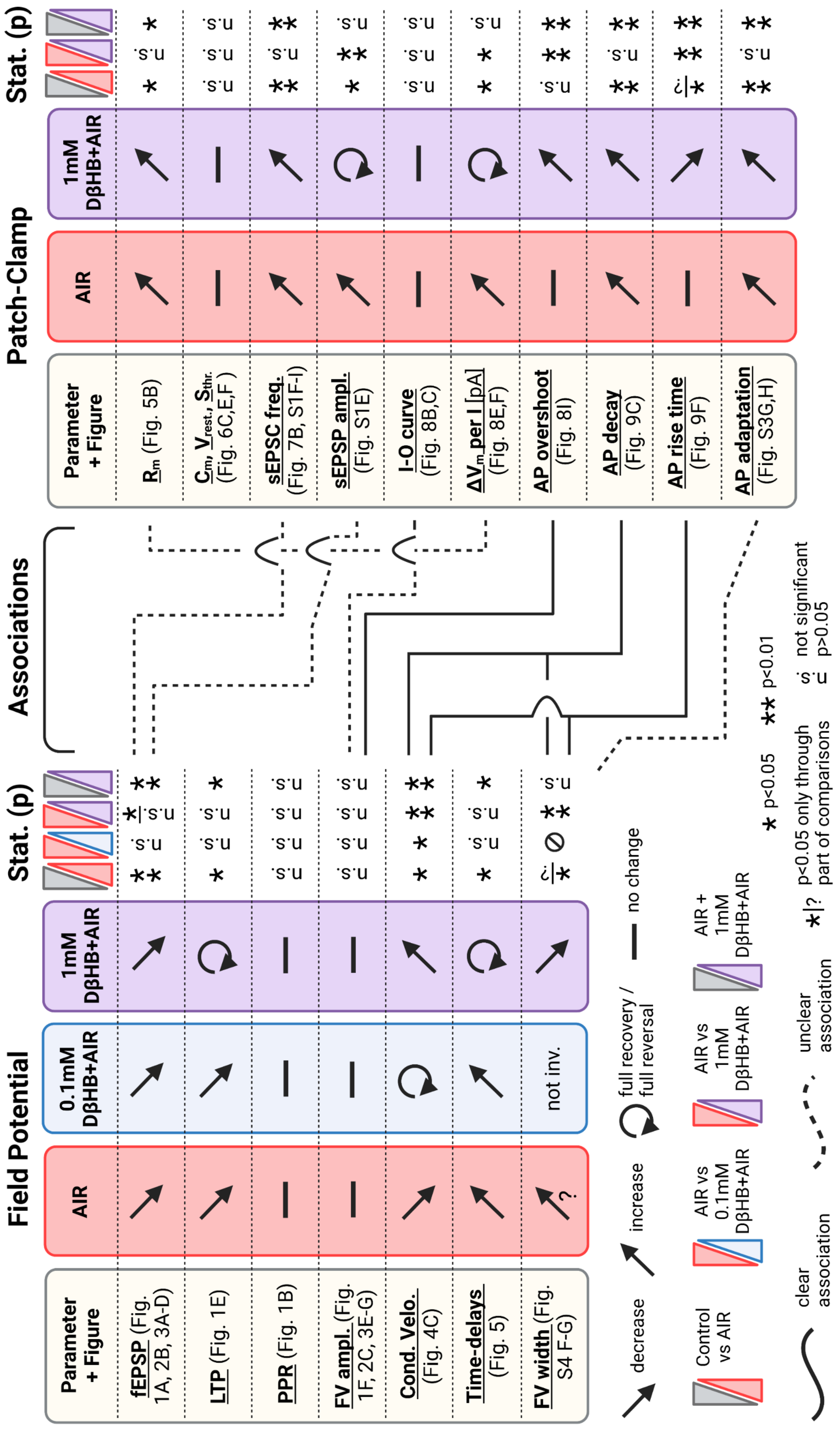

Field Potential
Patch-Clamp
Associations
Stat. (p)
Parameter + Figure
AIR
0.1mM DβHB+AIR
1mM DβHB+AIR
fEPSP (Fig. 1A, 2B, 3A-D)
LTP (Fig. 1E)
PPR (Fig. 1B)
FV ampl. (Fig. 1F, 2C, 3E-G)
Cond. Velo. (Fig. 4C)
Time-delays (Fig. 5)
FV width (Fig. S4 F-G)
not inv.
Rm (Fig. 5B)
Cm, Vrest, Sthr. (Fig. 6C,E,F )
sEPSC freq. (Fig. 7B, S1F-I)
sEPSP ampl. (Fig. S1E)
I-O curve (Fig. 8B,C)
ΔVm per I [pA] (Fig. 8E,F)
AP overshoot (Fig. 8I)
AP decay (Fig. 9C)
AP rise time (Fig. 9F)
AP adaptation (Fig. S3G,H)
decrease
increase
full recovery / full reversal
no change
Control vs AIR
AIR vs 0.1mM DβHB+AIR
AIR vs 1mM DβHB+AIR
AIR + 1mM DβHB+AIR
clear association
unclear association
*|? p<0.05 only through part of comparisons
* p<0.05
** p<0.01
n.s. not significant p>0.05

**Figure S8: Overall summary of key findings.**

Left: Most important results of the field potential recordings. Right: Most important results of the patch-clamp experiments. The black solid lines link field potential results with their counterparts during patch-clamp experiments. The dashed lines link field potential results with their most likely patch-clamp counterparts, although the associations are less clear. Figure prepared with BioRender.

Supplementary Methods

1.1 *Field potential recordings – data visualization. [Figure 3, Figure S2, Figure S5]*

To construct "heatmap" visualizations of the data, numerical matrices of the FV or fEPSP amplitude means were built for each of the four treatment groups. In every matrix, each new row represented responses during each subsequent train. Then, two additional columns were added at the end as a color normalization in the heatmap (0, 1 mV), and the matrix was saved as a text file. Next, the text files were imported as text images to ImageJ (NIH, USA), converted from 32- to 16-bit, saved as TIFF images and imported into Igor Pro (Wavemetrics, USA); then, their look-up table (LUT) was changed from "Grays" to "Spectrum."

The heatmaps for latency changes during train stimulation were generated similarly but using different normalizations: the 20 pulse train recordings had normalization boundaries of -1 and +1 ms.

1.2 *Multi-Gaussian fitting. [Figure S3]*

All data acquisition was performed using pCLAMP software (Molecular Devices, USA).

1.4 *Model of conduction velocity. [Figure 2]*

Here, we model how the AP shape affects the conduction velocity (CV) along an unmyelinated axon. We model an infinitesimally small unit of the axon consisting of three distinct units in a row, separated by a space h. The first and last of these units are kept "fixed" at voltages $V_1 = V_a$ (the peak membrane potential) and $V_3 = V_{rest}$, respectively; this potential difference leads to a voltage flow across the axon, which is generally described by the cable equation [68]:

$$\lambda^2 \frac{\partial^2 V}{\partial x^2} = \tau \frac{\partial V}{\partial t} + V$$

Once the intermediate unit $V_2$ is excited above a threshold voltage $V_t$, the unit is activated and self-excited up to a voltage $V_a$. This process then repeats *ad infinitum* along the entirety of the axon (assuming no backflow of the charge). The conduction velocity (i.e., the rate of this voltage flow) is thus inversely proportional to the reset time $t_{reset}$ (the first time when $V_2(t) = V_t$ and $V_2(0) = V_{rest}$). Over the discrete lattice described above, the cable equation is given by:

$$\lambda^2 \frac{V_3 - 2V_2 + V_1}{h^2} - \tau \frac{dV_2}{dt} = V_2$$

Solving this for $V_2(t)$ and substituting in the boundary conditions, we find:

$$V_2(t) = \frac{\lambda^2 (V_{rest} + V_a)}{h^2 + 2\lambda^2} + K e^{\frac{-1}{\tau}\left(1 + \frac{2\lambda^2}{h^2}\right)t} = V_B + K e^{\frac{-1}{\tau}\left(1 + \frac{2\lambda^2}{h^2}\right)t}$$

where $K$ is an integration constant needed to satisfy the initial condition $V_2(0) = V_{rest}$. Substituting in the initial condition and solving for $t_{reset}$, we find:

$$t_{reset} = \frac{-\tau}{1 + \frac{2\lambda^2}{h^2}} \log \frac{V_t - V_B}{V_{rest} - V_B}$$

Thus:

$$CV \propto \frac{1}{\log \frac{V_{rest} - V_B}{V_t - V_B}}$$

However, h is an arbitrarily small spacing that can approach zero. Thus:

$$CV \propto \frac{1}{\log \frac{V_{rest} - V_a}{2V_t - V_{rest} - V_a}}$$

1.5 *Hodgkin-Huxley model. [Figure S7]*

The sample sizes for all statistical comparisons in this work were determined based on the means and pooled standard deviations from preliminary recordings of 8-10 cells or slices per group, $\alpha$ = 0.05, $\beta$ = 0.8, corrected for the number of pairwise comparisons ($\tau$), based on the following equations:

$$n=2\left(\sigma\frac{z_{1-\alpha/(2\tau)}z_{1-\beta}}{\mu A-\mu B}\right)^2,$$

$$1-\beta=\phi\left(z-z_{1-\alpha/(2\tau)}\right)+\phi\left(-z-z_{1-\alpha/(2\tau)}\right),$$

$$z=\frac{\mu A-\mu B}{\sigma\sqrt{\frac{2}{n}}},$$

where n is the sample size, σ is the standard deviation, $\phi$ is the standard normal distribution function, $\alpha$ is the Type I error rate, $\tau$ is the number of pairwise comparisons, and $\beta$ is the Type II error rate. During the calculations, the normality of the residuals and equality of the variances were assumed *a priori*.

All of the calculations were performed with an online calculator available at:

http://powerandsamplesize.com/Calculators/Compare-k-Means/1-Way-ANOVA-Pairwise
